# Promoter strength and position govern promoter competition via transcript-dependent insulation

**DOI:** 10.1101/2025.05.06.652547

**Authors:** Mervenaz Koska, Masahiro Nagano, Tomek Swigut, Alistair Nicol Boettiger, Anders S. Hansen, Joanna Wysocka

## Abstract

Long-range competition between promoters within a shared regulatory landscape has been implicated in development and disease, but the determinants of promoter competition remain unclear. Here, we introduce diverse promoters into defined genomic sites within the *Sox2* locus and measure how these insertions attenuate endogenous *Sox2* expression. We find that the level of reduction in endogenous *Sox2* transcription is correlated with the strength of the inserted promoter. Transcription from the inserted promoter is required for competition, with longer transcripts resulting in more competition. The inserted active promoter and its associated transcriptional unit function as an insulator, rendering competition position-dependent. Competition is counteracted by the HUSH-mediated silencing of the inserted promoters. Together, our work uncovers the rules governing promoter competition, highlights its impact on tuning gene expression levels and genome evolution, and suggests that transcriptional units producing transcripts of sufficient level and length can mediate insulation independently of CTCF and cohesin.

## INTRODUCTION

Gene expression is coordinated through an interplay between multiple classes of cis-regulatory elements (CREs), such as enhancers, promoters, and boundary elements^1–4^. Typically, multiple enhancers and promoters are present within a given regulatory domain, raising the question of how CREs influence each other’s activity^5,6^. One previously described phenomenon is ‘promoter competition’, whereby a promoter can dampen the transcriptional output of another promoter located in cis^7–9^. While promoter competition can occur through transcriptional interference at closely spaced genes^10–13^ it has also been observed over long distances that cannot be explained by interference. A canonical example is the beta-globin locus, where the fetal and adult globin promoters compete for the distal locus control region^7,14^. In this system, activation of the adult b-globin promoters facilitates shutdown of the fetal globin promoter through competition^7^. Conversely, disrupting any of the adult b-globin promoters redirects the b-globin enhancers to fetal globin promoter and reactivates its expression^15^.

Although promoter competition likely occurs at many loci across development and disease, such effects can be challenging to reveal without perturbing a given promoter and assaying the impact on other promoters in cis. Promoter competition is even more noticeable when a *de novo* promoter is introduced into a genomic region and alters the activity of the endogenous promoters nearby. A compelling example is the disease-causing single nucleotide variant in the human alpha-globin locus that generates a transcription factor binding site, inducing transcription^16^. This *de novo* promoter disrupts the communication between the a-globin enhancer cluster and its cognate promoters, thus leading to reduced a-globin gene expression and causing a-thalassemia^16^.

While these examples suggest that promoter competition broadly shapes gene expression, the rules governing this phenomenon have not been systematically studied. We therefore explored how a cis-regulatory network responds to the introduction of a new promoter at the well-studied *Sox2* locus in mouse embryonic stem cells (mESC), where the majority of *Sox2* expression is controlled by a super enhancer (SE) that lies ∼110 kb away from the promoter^17,18^. We found that attenuation of endogenous *Sox2* promoter activity correlates inversely with inserted promoter’s strength, with only the strongest promoters showing substantial competition. Moreover, competition also depends on the position of the inserted promoter in relation to the endogenous promoter and SE, with the inserted promoter acting as an insulator, reducing contacts between the SE and *Sox2.* This insulation-mediated competition does not require CTCF or cohesin, and instead depends on transcription from the inserted promoter, with longer transcripts resulting in stronger competition. Lastly, promoter competition is counteracted by the Human Silencing Hub (HUSH)^19,20^, whose loss exacerbates promoter competition. Altogether, our work shows that promoter strength and position – and the related level and length of the resulting transcripts – are key determinants of promoter competition.

## RESULTS

### A synthetic platform to study determinants of promoter competition

To systematically introduce a new promoter at the *Sox2* locus, we used a previously generated Cast/129 hybrid mESC line, with the *Sox2* gene tagged with P2A-eGFP on the Cast allele and P2A-mCherry on the 129 allele^21^. To enable controlled insertion at a defined location, we leveraged the landing pad with heterospecific Flippase Recognition Sites (FRT and FRT3) between *Sox2* and its SE on the eGFP/Cast allele (Fig.1a)^21^. To target the landing pad, we generated donor plasmids with matching heterospecific FRT sites flanking a 1kb promoter fragment (850 bp upstream and 150 bp downstream of the most prominent TSS), followed by the mTagBFP2 coding sequence with an intron, and an SV40 polyA signal (Fig.1a, Supplementary Table 1). We clonally integrated a small library of housekeeping and developmental promoters, spanning diverse transcription factor and cofactor binding profiles and activities in their endogenous context (Fig. 1b, Extended Data Fig. 1a). Additionally, we included positive control sequences corresponding to the strong promoters Ef1a and CAG, and negative controls, with either no promoter insert or an E. coli coding sequence (CDS) lacking regulatory activity in eukaryotic cells (hereafter referred to as the ‘neutral’ sequence) (Fig. 1b). All sequences were integrated in forward or reverse orientation (Fig. 1a), and multiple clonal mESC lines were isolated for each insert. This synthetic platform allows simultaneous measurement of ectopic promoter activity (via mTagBFP2), endogenous *Sox2* promoter activity in cis from the insertion (via eGFP) and activity of the unperturbed *Sox2* allele without the landing pad (via mCherry) as a reference. Under promoter competition, we expect the eGFP levels to decrease in the presence of the mTagBFP2, whereas mCherry should be unaffected (Fig. 1c).

**Figure 1.**
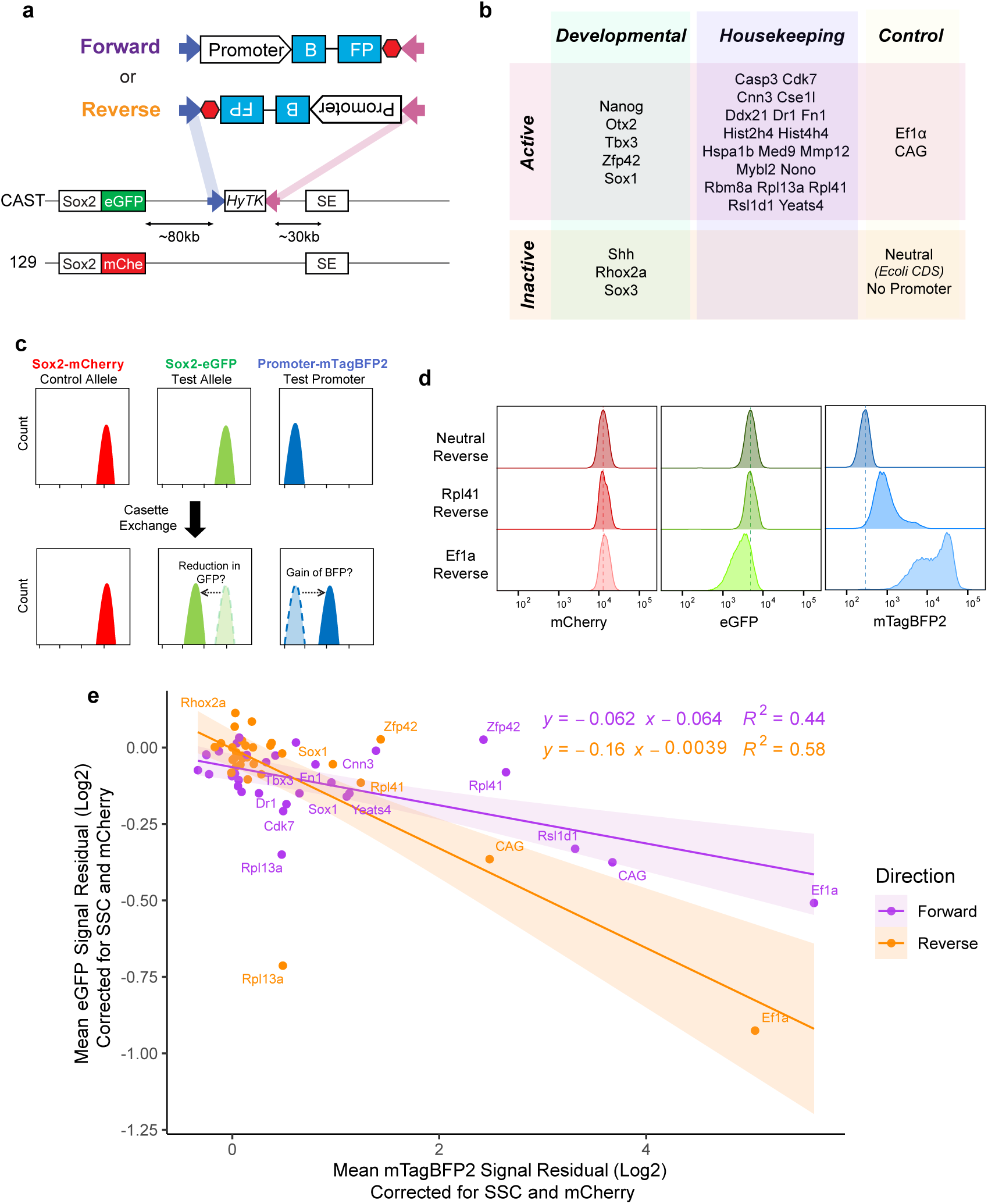
Promoter competition screen at the Sox2 locus. **a)** Schematic of Sox2 locus with the landing pad with hygromycin phosphotransferase-thymidine kinase fusion gene HyTK, flanked by heterospecific FRT sites, and of the inserts that are integrated at those sites either in the forward or in reverse orientation. Blue arrow represents FRT, pink arrow FRT3. SE, Super-enhancer. **b)** List of promoters tested. **c)** Mock flow cytometry histogram data demonstrating expected changes in fluorescent reporter signal in the competition scenario. **d)** Sample flow cytometry data from three integrations, representing neutral sequence and two tested promoters. Dashed lines are at the median of neutral insert. **e)** Scatterplot of mean corrected eGFP and mTagBFP2 signal from all promoters. Each dot is the average value from 3 biological replicates (measurements from 3 independent clonal lines), each measured in triplicate. The values are the mean of the residual for fluorescence intensity after correcting for SSC and mCherry (see methods). Equation and R^2^ value for linear regression for each direction is displayed. Values translated vertically and horizontally to make neutral insert adjusted to 0.

### Promoter competition is dependent on the strength of the inserted promoter

To assess promoter competition, we measured mCherry, eGFP and mTagBFP2 fluorescence by flow cytometry in three clonal lines per promoter insert in both forward and reverse directions. As expected, mCherry remained stable across cell lines (Extended Data Fig. 1b). We nonetheless performed a linear regression correction on eGFP and mTagBFP2 measurements for SSC and mCherry to account for small variations that might arise due to cell size and state (See Methods). We next plotted the corrected mean fluorescence intensity for eGFP and mTagBFP2 (Fig. 1e) and observed an inverse correlation. That is, with increasing mTagBFP2 levels from the inserted promoter, the expression driven by the endogenous *Sox2* promoter was progressively reduced, suggesting that promoter strength (approximated here by mTagBFP2 expression) is one of the determinants of competition (Fig. 1d,e). Interestingly, the extent of the competition was also dependent on the orientation of the promoter insert, as demonstrated by differences in slopes of linear regression fits for forward and reverse integrations, with reverse integrations showing more competition (Fig. 1e); we elaborate on this observation when discussing Figure 7.

### Promoter competition can be modulated by CTCF sites in the promoter

The Rpl13a promoter in the reverse orientation was a strong outlier to the trendline, with noticeably reduced *Sox2* eGFP despite moderate mTagBFP2 levels (Fig. 1e). To investigate, we examined the genomic features of the Rpl13a promoter and identified three tandem CTCF binding motifs oriented in the same direction (Extended Data Fig. 1c). We reasoned that these CTCF sites may disrupt SE-*Sox2* promoter communication by creating a boundary that blocks cohesin-mediated loop extrusion (Extended Data Fig. 1d). Indeed, previous work showed that tandem CTCF sites convergent with those at the SE can attenuate *Sox2* expression^21^. Such convergent orientation is present in the reverse Rpl13a promoter insert (Extended Data Fig. 1d). To test whether CTCF sites contribute to the Rpl13a outlier effect, we integrated mutant versions of the Rpl13a promoter with either deleted or scrambled the CTCF motifs. In forward Rpl13a integration, the mutant motifs had a negligible effect on both mTagBFP2 and eGFP signal relative to unmodified Rpl13a. Yet in reverse direction, deletion or scramble of CTCF motifs both led to decrease of mTagBFP2 and rescued the eGFP reduction (Extended Data Fig. 1e). Accordingly, ChIP–qPCR confirmed that CTCF occupancy at the Rpl13a promoter was reduced by ∼30% in both directions, as expected given the presence of two intact endogenous Rpl13a alleles in addition to the integration (Extended Data Fig. 1f). These data argue that presence and orientation of CTCF sites at the promoters can influence promoter competition.

### Allelic series of Ef1a promoter augments correlation between promoter strength and competition

To test whether promoter strength drives competition, we synthesized Ef1a deletion mutants that varied in strength (Fig. 2a) based on previous work^22^. These mutants spanned over an order of magnitude range in activity in an episomal luciferase assay, confirming effective modulation of promoter strength (Fig. 2b). When these mutant Ef1a promoters were integrated clonally into our landing pad, we once again detected an inverse correlation between eGFP and mTagBFP2 levels, suggesting that competition is driven by promoter strength (Fig. 2c). To confirm that the competition reflects intrinsic promoter strength rather than differential compatibility with the SE, we also independently measured the activity of the remaining endogenous promoters with luciferase assay (Extended Data Fig. 2a). Overall, luciferase activity and BFP levels from the integrated reporters were strongly correlated (Extended Data Fig. 2b), and this relationship held for both constitutive promoters (such as Ef1a and its mutants, Rpl41, and CAG) and developmental promoters (such as Zfp42, Nanog, Sox1 and Sox3), although the latter were generally weaker (Extended Data Fig. 2a). In line with its low intrinsic activity, even the Sox2 promoter showed minimal BFP and negligible effects on eGFP when integrated at the landing pad (Extended Data Fig. 2e, f). Together, these results suggest that the inherent promoter strength is the major determinant of promoter competition, regardless of the ‘compatibility’ of the promoter with the enhancer. Combining all tested promoters further illustrates the relationship between promoter strength and competitive effects over a broad range of promoter types and activities (Extended Data Fig. 2c, d).

**Figure 2.**
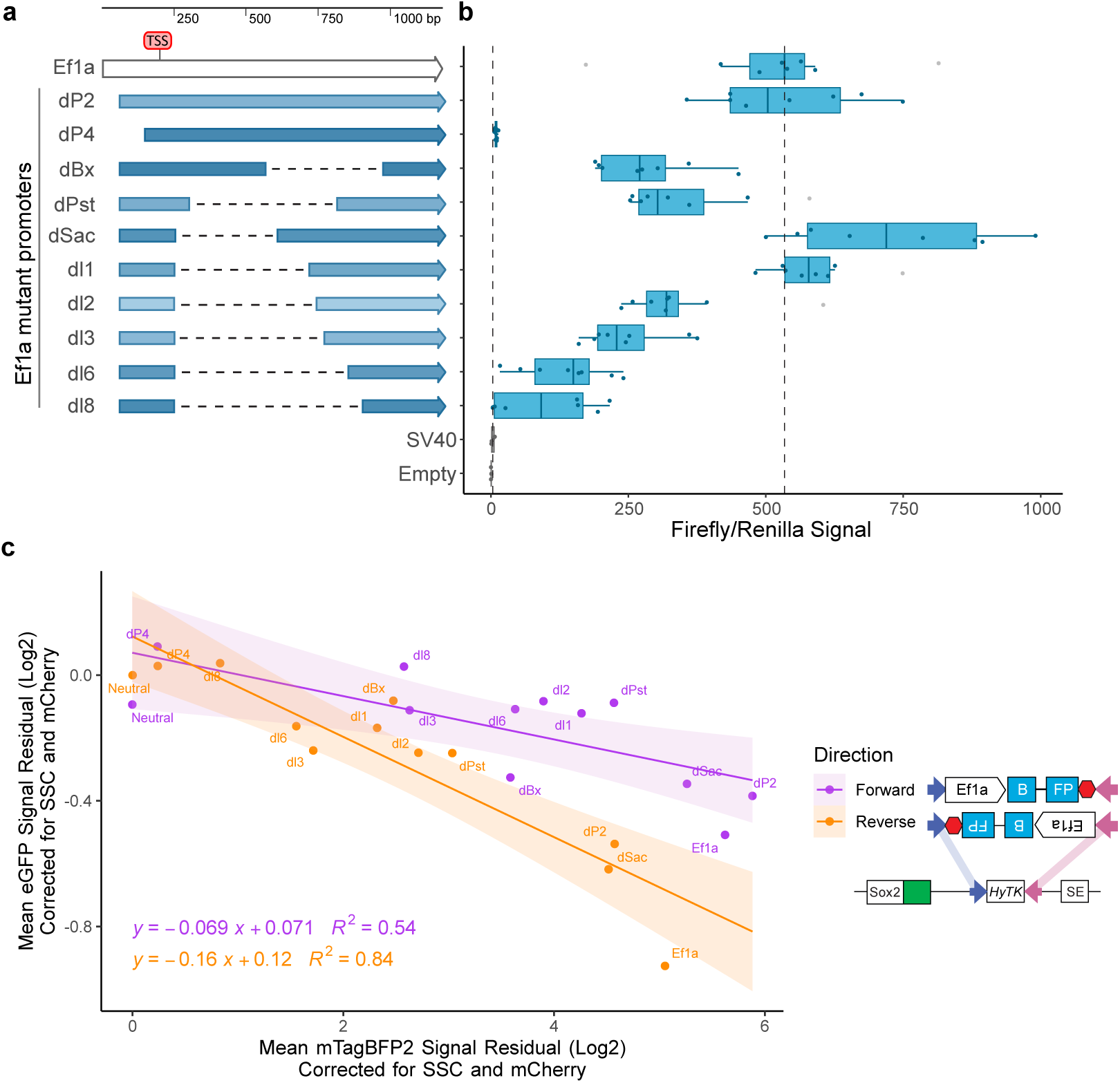
Allelic series of Ef1a promoter recapitulates inverse correlation between expression from integrated and endogenous promoters. **a)** Schematic of allelic series of Ef1a promoter, dashed lines signify deleted regions. **b)** Episomal Luciferase reporter assay measurements from each mutant promoter. Measurements performed twice with two distinct preps of reporter plasmid, transfected in duplicate. Firefly divided by Renilla signal is displayed. Boxes represent the median, IQR, and whiskers (±1.5 IQR), outlier measurements displayed in gray. Boxplots for control measurements with empty vector and SV40 promoter are filled with gray. Dotted line represents the median of the SV40 and Ef1a measurements. **c)** Scatterplot of mean corrected eGFP and mTagBFP2 signal of mutant Ef1a promoters for 3 biological replicates (independent clonal cell lines), measured in triplicate. Equation and R^2^ value for linear regression for each direction is displayed. Values translated vertically and horizontally to make neutral insert adjusted to 0. Schematics of integration in forward and reverse orientation is shown in the inset.

### Loss of HUSH-mediated silencing enhances promoter competition

We observed that many promoters displayed a bimodal mTagBFP2 expression pattern, with high-and low-expressing populations (Fig. 3a), reminiscent of variegated transgene silencing by the Human Silencing Hub Complex (HUSH)^19,23^. HUSH is a transcription-dependent silencing complex that targets non-self DNA such as transgenes or transposable elements^23^ by depositing H3K9me3 to attenuate their expression^23^. This was surprising, given that we have included an intron in the mTagBFP2 coding sequence, which was shown to counteract HUSH silencing^24^. However, consistent with silencing, ChIP-qPCR analysis at the Ef1a promoter or the neutral (non-expressing) integrations detected H3K9me3 enrichment at the integration sites, with a substantially higher H3K9me3 deposition at the Ef1a reporter compared to a neutral sequence (Fig. 3b,c). Because establishment of HUSH silencing requires transcription^20,25^, we hypothesized that increased H3K9me3 at the active Ef1a reporter underlies the bimodal mTagBFP2 signal. Consistent with this, knockout of the core HUSH subunit MPP8 ((Extended Data Fig. 3a). abolished the low mTagBFP2 population (Fig. 3d, compared to 3a, and Fig. 3e) and eliminated H3K9me3 at the Ef1a reporter (Extended Data Fig 3d). Similar loss of bimodality was observed for reporters driven by other promoters as well, such as CAG and Rpl41 (Fig. 3d). Thus, even in the presence of an intron and a strong promoter, the inserted reporters are susceptible to HUSH-mediated silencing.

**Figure 3.**
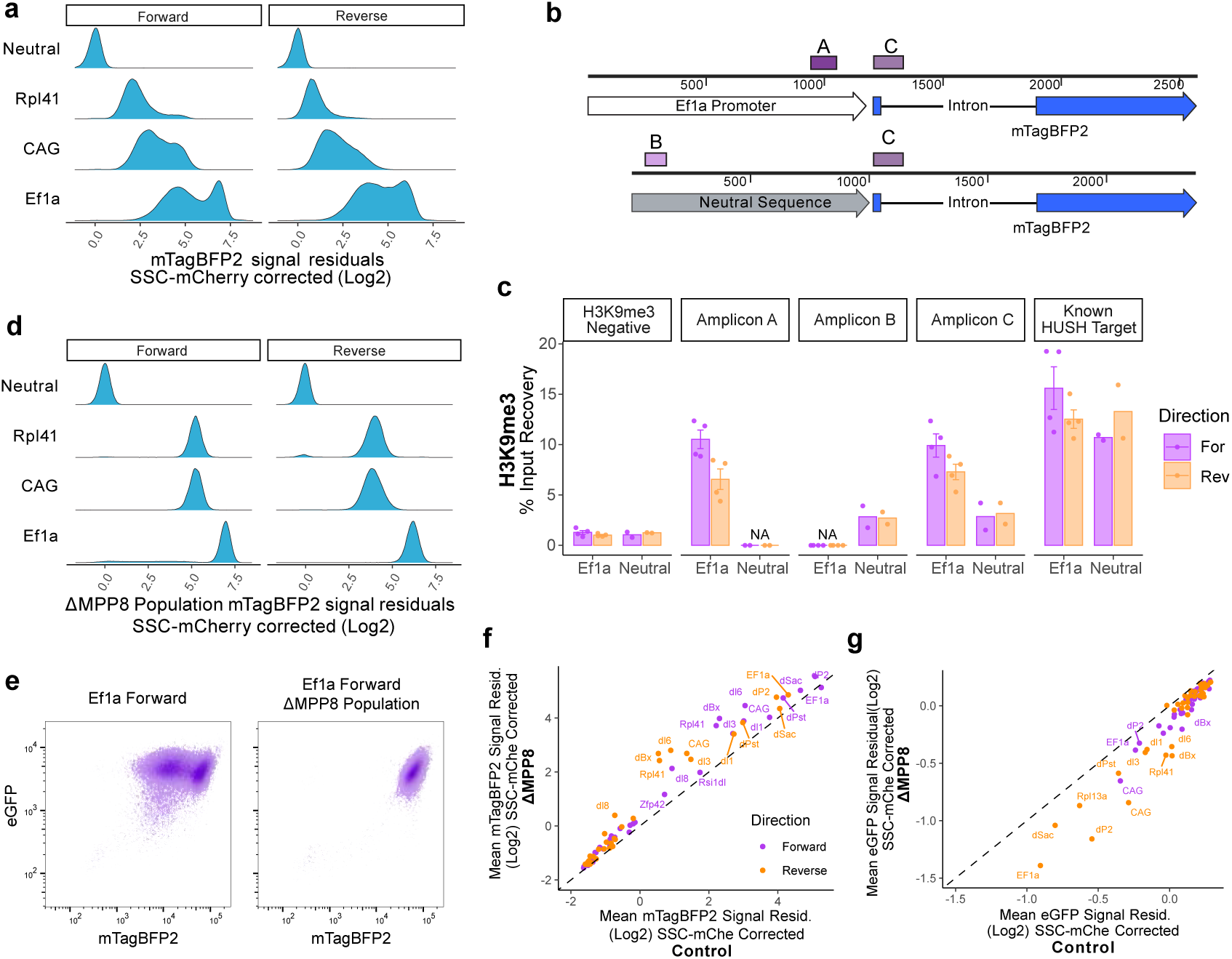
HUSH mediated silencing at promoter inserts. **a)** Density plot of mTagBFP2 residual after correction for SSC and mCherry for a subset of promoters integrated in the WT background. **b)** Schematic of inserts with location of ChIP-qPCR amplicons indicated with purple bars. Each of the inserted promoter sequences have a unique amplicon (Ef1a Amplicon A or neutral sequence Amplicon B), and an amplicon corresponding to the exon-intron junction of the inserted mTagBFP2 reporter (shared between all cell lines, Amplicon C) in addition to H3K9me3-negative and positive regions outside of the Sox2 locus, which are not depicted here. **c)** H3K9me3 ChIP-qPCR percent input recovery at amplicon sites labeled in (B). n=4 for Ef1a, n=2 for Neutral inserts, bars represent mean; points represent individual measurements, ± s.e displayed for Ef1a. **d)** Density plot of mTagBFP2 residual after correction for SSC and mCherry for a subset of promoters in ΔMPP8 populations. **e)** Raw flow measurements of eGFP and mTagBFP2 signal in Ef1a Forward insert in WT and ΔMPP8 backgrounds. **f)** Scatterplot comparing mean corrected mTagBFP2 signal in ΔMPP8 and control edited populations. Dashed line, y=x. **g)** Scatterplot comparing mean corrected eGFP signal in ΔMPP8 and control edited populations. Dashed line, y=x.

We next examined how loss of HUSH affects promoter competition by knocking out MPP8 at a population level in all our promoter insert lines (Extended Data 3a). We saw that HUSH removal did not change the inverse correlation between eGFP and mTagBFP2 (Extended Data 3b). As expected, loss of MPP8 yielded an increase in the mean mTagBFP2 expression from all promoters relative to nontargeting guide-treated control cells (Fig. 3f). Conversely, expression of eGFP driven by the *Sox2* promoter was decreased across all ΔMPP8 cells compared to controls (Fig. 3g). Taken together, in the absence of HUSH-mediated silencing, we observe elevated expression from the inserted promoters and increased competition with the *Sox2* promoter.

### Promoter competition is position dependent

Next, we asked whether the competition depends on the location of the inserted promoter between the endogenous promoter and the SE. We used an alternative version of the parental cell line, which has the landing pad at an equidistant location (∼30kb away) but downstream from the SE (Fig. 4a)^21^. We integrated a subset of the promoters at this downstream landing pad and compared their activity to their in-between landing pad counterparts. As expected, downstream insertions showed a range of mTagBFP2 expression (Fig. 4b). However, unlike in-between insertions, they had no effect on the *Sox2* eGFP expression (Fig. 4b), indicating that promoter competition is indeed dependent on the genomic location. Of note, downstream integrations often showed higher mTagBFP2 expression than their in-between counterparts, which may suggest a reciprocal competition between *Sox2* and the integrated promoter.

**Figure 4.**
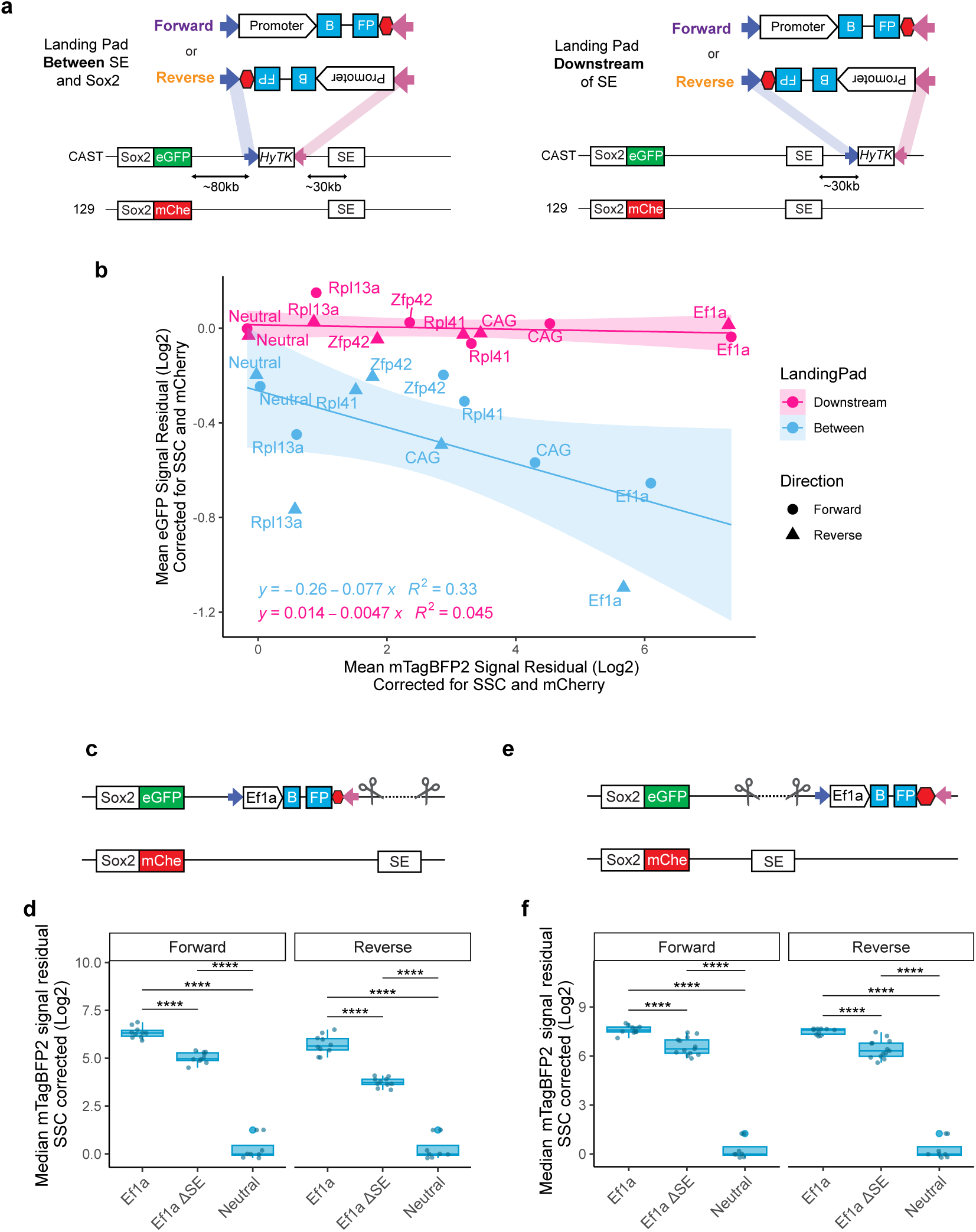
Promoter competition is position dependent. **a)** Schematic comparing two landing pad locations: between SE and Sox2 gene (left) and downstream of the SE (right). **b)** Scatterplot of corrected mean eGFP and mTagBFP2 signal for subset of promoters tested in both at the “between” and “downstream” landing pads. Mean of 3 biological replicates, measured in triplicate. Values translated vertically and horizontally to make neutral insert adjusted to 0. Equation and R^2^ value for linear regression for each landing pad is displayed. **c)** Schematic of allele-specific SE deletions in cell lines with the Ef1a promoter integration at the “between” landing pad. **d)** Corrected median mTagBFP2 measurements from in between landing pad lines with Ef1a integrated, Ef1a integrated with ΔSE and neutral insert controls. Median from 3 biological replicates (independent clonal lines), measured in triplicates plotted together. Boxes represent the median, IQR, and whiskers (±1.5 IQR). Pairwise t-tests with Bonferroni correction, ****Padj ≤ 0.0001. **e)** Schematic of allele specific SE deletions in cell lines with the Ef1a promoter integration at the “downstream” landing pad. **f)** Corrected median mTagBFP2 measurements from downstream landing pad lines with Ef1a integrated, Ef1a integrated with ΔSE and neutral insert controls. Median from 3 biological replicates (independent clonal lines), measured in triplicates plotted together. Boxes represent the median, IQR, and whiskers (±1.5 IQR), large blue dot marks outlier. Two-sided pairwise t-tests with Bonferroni correction, ****Padj ≤ 0.0001. In **d** and **f** values translated vertically to make neutral insert adjusted to 0.

Although the downstream landing pad is within the same TAD as *Sox2* and SE^26^, it lies outside of the *Sox2*-SE loop demarcated by the convergent CTCF sites^27,28^(Extended Data Fig 4b). We therefore considered whether the lack of competition from promoters inserted at this site might reflect their failure to receive regulatory input from the SE. To test this, we deleted the SE at the Cast allele in cell lines with Ef1a integrated at either the in-between or downstream landing pad (Fig. 4c, e). As expected, SE deletion abolished most eGFP expression across all cell lines (Extended Data Fig. 4a). Interestingly, while the median mTagBFP2 expression was overall higher at the downstream landing pad location, SE deletion impacted the mTagBFP2 levels regardless of location (Fig. 4d, f). These data indicate that despite strong inherent activity, expression from the Ef1a promoter is comparably boosted by the SE in both locations. Consistent with the SE’s ability to boost expression of Ef1a promoter regardless of its position, mutation of the CTCF sites within the SE did not substantially alter the position-dependent competition, as might have been expected if these CTCF sites strongly influenced directionality of the SE contacts (Extended Data Fig 4b, c). Instead, we observed no effect of these mutations on eGFP levels when the reporter was inserted in the in-between landing pad, whereas in the downstream location, we detected a small reduction in eGFP, commensurate with previously reported effect of these CTCF sites deletion on *Sox2* expression^28^. Taken together, these results argue against a simple enhancer resource sharing model underlying promoter competition and instead emphasize the importance of relative promoter position.

### Active reporter insertions create a boundary, with insulation correlated with competition

Given the privileged role of the in-between location for promoter competition, we explored whether active insertions act as a boundary that attenuates enhancer-promoter communication. To investigate, we performed Region Capture Micro-C (RCMC) at the *Sox2* locus^27^ in Ef1a, Rpl13a, and neutral insert lines in both orientations. We used the Rpl13a Reverse promoter as a positive control, as its three CTCF sites^21^ in this orientation are predicted to loop with the SE, creating a bona fide boundary^29–31^ for the native *Sox2* promoter (Extended Fig. 1d).

Leveraging the SNPs in the hybrid cell lines we generated allele-specific contact maps^32^ (see Methods). As expected, in each examined cell line, the reference 129 allele without the insert showed a strong dot corresponding to the *Sox2-*SE contacts (Fig 5a-d, Extended Data Fig 5a,b). On the Cast allele, this dot was also evident in neutral control lines (Fig 5c,d), and to a lesser extent, in the Rpl13a Forward inserts (Extended Data Fig 5a). In contrast, the Rpl13a Reverse (Extended Data Fig 5b) and Ef1a insert in both orientations showed weakening of the *Sox2-*SE dot (Fig 5a,b), and emergence of an insulating boundary at the insertion site. To quantify, we calculated insulation scores ^33,34^ for each insert and observed that Ef1a in the reverse orientation shows stronger insulation than the bona fide boundary formed by the three CTCF sites at the Rpl13a promoter (Extended Fig 5c). Across all insertions, reduction in Sox2-eGFP expression strongly correlated with insulation strength (Fig. 5e) and with decreased Sox2–SE contact probability (Fig. 5f). However, there was no correlation between SE-insert contact probabilities and mTagBFP2 expression, consistent with the majority of the mTagBFP2 expression being driven by the intrinsic activity of the promoter (Fig. 5g). Overall, our results support a model in which competing promoters form an insulating boundary that reduces enhancer-promoter loop strength and gene expression levels.

**Figure 5.**
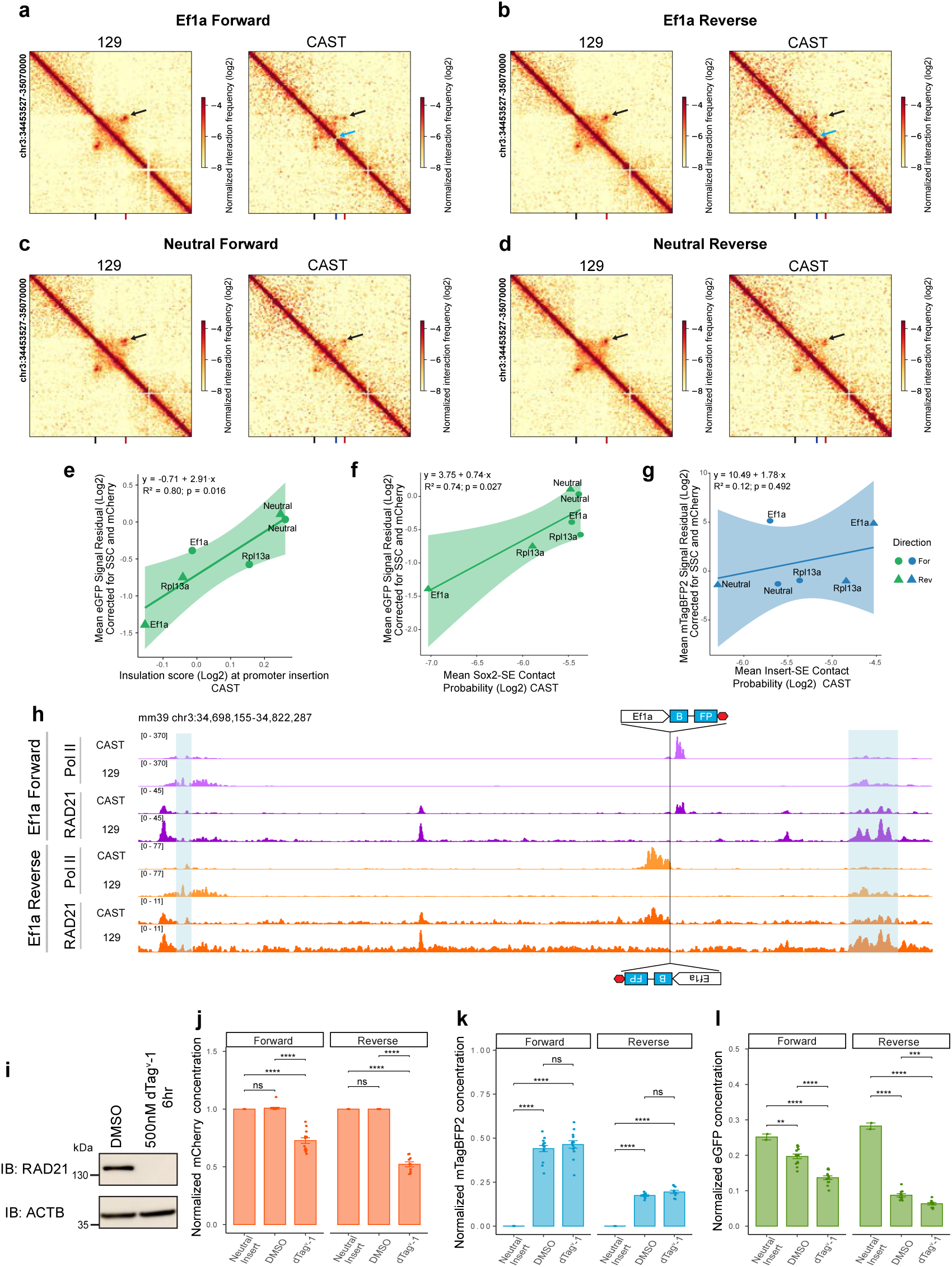
Active promoter inserts change the conformation of the locus, but competition does not require cohesin. a,b,c,d) Contact map visualization of RCMC data for Ef1a For (a), Ef1a Rev (b), Neutral For (c) and Neutral Rev (d) integrations at 5-kb resolution. Contacts map for control allele 129 shown on the left, for insert allele CAST on the right. Sox2 location is marked with a black notch, insert with blue, and SE with red. Black arrows highlight SE-Sox2 contact, blue arrows highlight domain split. Maps generated with merged reads from two biological replicates (independent clones). **e)** Scatterplot of corrected mean eGFP signal and insulation scores (lower score means higher insulation) at the promoter insert. **f)** Scatterplot of corrected mean eGFP signal and Sox2-SE contact probability. **g)** Scatterplot of corrected mean mTagBFP2 signal and insert-SE contact probability. **e,f,g** Shape legend shown at the inset of **g**. Equation, R^2^ and p-value for linear regression is displayed for each plot. **h)** Allele specific RNA Pol II and RAD21 ChIP-seq tracks in Ef1a integration cell lines. Landing pad location and direction marked with cartoon of Ef1a integration, blue highlights mark Sox2 coding sequence and SE. **i)** RAD21 western blot in representative Rad21-FKBP12^F36V^ clonal line in DMSO control and after 6 hr depletion. Membranes were probed with antibodies against RAD21 and ACTB as a loading control. Representative of two independent experiments. **j,k,l)** RT-qPCR measurements of **(j)** mCherry, **(k)** mTagBFP2, **(l)**eGFP transcripts after 6 hrs of RAD21 depletion. Absolute concentrations were first normalized to housekeeping gene Gapdh, then normalized to the corresponding mCherry control. 6 clonal cell lines were assayed for each direction, in duplicate, mean ± s.e. Two-sided pairwise t-tests with Bonferroni correction, **Padj ≤ 0.01; ***Padj ≤ 0.001; ****Padj ≤ 0.0001.

### Cohesin accumulates at the active reporter but is not required for competition

Having established that active promoter integrations alter the 3D landscape of the locus we next asked how this effect was mediated. While the Ef1a promoter lacks CTCF binding, RNA Polymerase II (Pol II) has also been proposed to act as a barrier to cohesin mediated loop extrusion ^35–39^. To investigate insert-mediated changes in the localization of Pol II and RAD21, a cohesin subunit, we performed ChIP-seq against Pol II and RAD21 in cell lines with Ef1a, Rpl41 and the inactive neutral reporter. To avoid confounding effects from HUSH silencing, we used ΔMPP8 cell lines (Extended Data Fig. 3c,d). In Ef1a integrated cell lines we detected a Pol II pile-up at the 3’ end of the reporter relative to the direction of transcription, consistent with transcriptional termination zone (Fig. 5h). Additionally, we observed RAD21 peaks that co-occur with Pol II (Fig. 5h) near these integrations. We saw a similar, albeit smaller in magnitude, allele-specific pile-up of RAD21 and Pol II at the region downstream from the Rpl41 reporter as well (Extended Fig. 5d). These peaks were not present on the 129 allele, nor at the Cast allele with neutral integrations, indicating that the accumulation of cohesin at this site depends on promoter activity (Extended Data Fig. 5d).

To test whether cohesin is required for promoter competition, we acutely depleted RAD21 using an FKBP12^F36V^ degron system^40^ in the Ef1a reporter lines (Fig. 5i) and measured *mCherry*, *eGFP* and *mTagBFP2* transcript levels using RT-qPCR (Fig. 5j-l). Loss of cohesin resulted in a decrease in *mCherry* levels, consistent with the previous reports indicating that cohesin loss decreases *Sox2* transcription by ∼25%^41^ (Fig. 5j), and it led to a modest (but statistically insignificant) increase in *mTagBFP2* expression (Fig. 5k). If loss of cohesin would alleviate promoter competition, then *eGFP* transcript levels should recover in dTag^v^-1-treated samples compared to the DMSO treated controls, even after accounting for the decreases associated with the effect of cohesin on the endogenous *Sox2* expression, as measured by the reduction in *mCherry*. However, the effects of cohesin depletion are comparable at the *mCherry* and the *eGFP* transcripts, with a reduction of *eGFP* expression in both orientations (Fig. 5l). This argues that cohesin is not necessary for competition and that active promoters can mediate insulation in a CTCF and cohesin-independent manner.

### Blocking transcription downstream from the TSS rescues promoter competition

Given the correlation between competition and promoter strength (i.e., transcript levels), we wondered if blocking transcription through the inserted reporter would alleviate competition. To test this, we employed our previously described CARGO approach.^42^ We assembled an array of six co-directional guides targeting the 5’ intron region of the *mTagBFP2* sequence (Gene-targeting, GT CARGO) (Fig. 6a) that will recruit dCas9, with a goal of physically blocking passage of Pol II through the gene without affecting assembly of transcriptional machinery at the promoter^43^. Once again, we used the ΔMPP8 Ef1a or neutral lines (Extended Data Fig. 3c) to avoid confounding effects of HUSH silencing.

**Figure 6.**
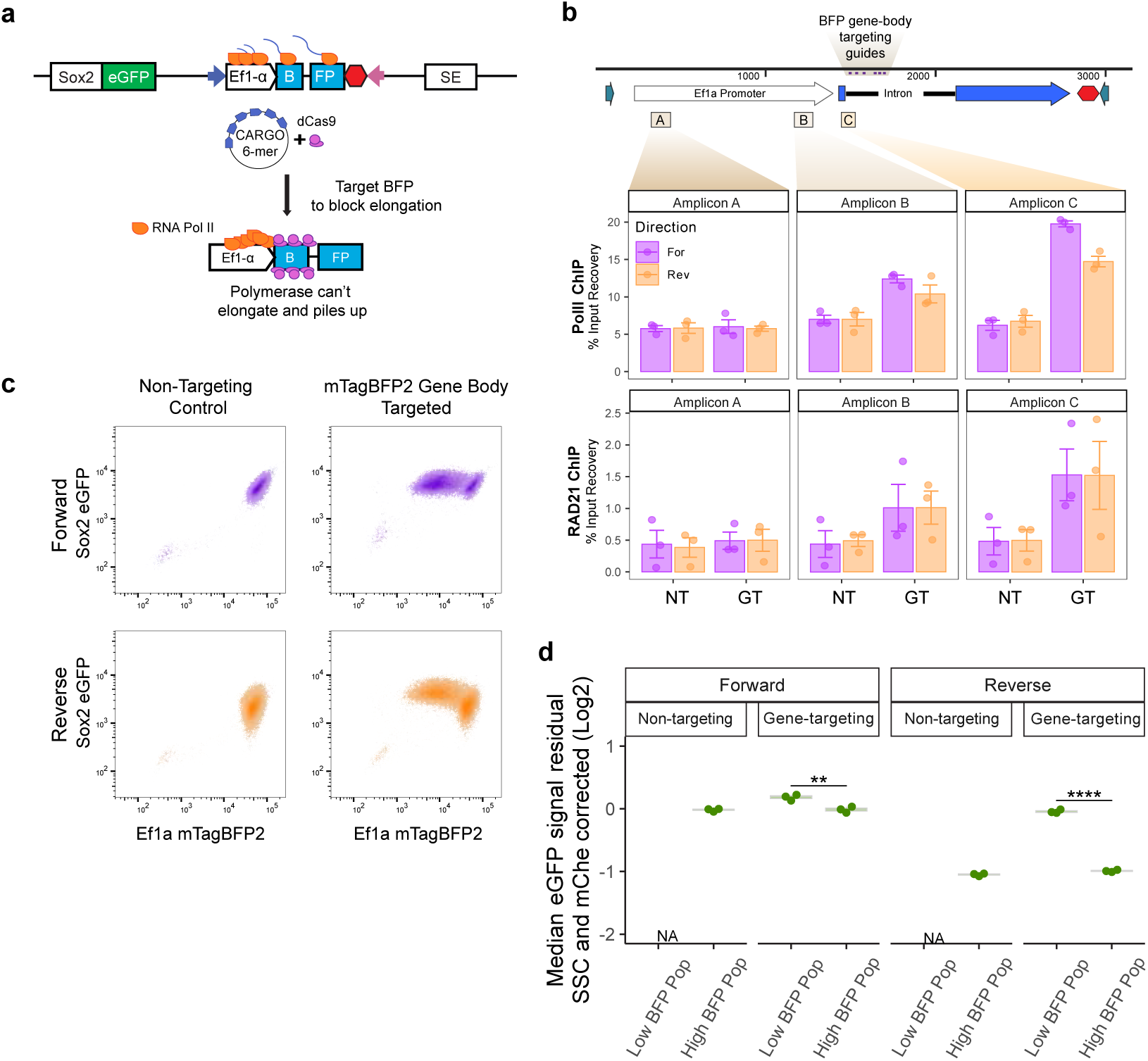
Blocking transcription with dCas9-CARGO rescues competition. **a)** Schematic of dCas9-CARGO strategy. A 6-mer CARGO guide RNA array targeting the gene body of mTagBFP2 and dCas9 is introduced to the cells with the goal of blocking elongation. **b)** RNA Pol II and RAD21 ChIP-qPCR percent input recovery at highlighted amplicon sites. n=3, mean ± s.e is plotted, NT=Non-targeting control, GT=Gene targeting CARGO. CARGO guides’ target locations are labeled with purple lines and highlighted in beige. **c)** Flow measurements of eGFP and mTagBFP2 signal in Ef1a promoter insert cells expressing dCas9 and either non-targeting control or gene-targeting CARGO array from a representative replicate. **d)** Corrected median eGFP measurements from “Low BFP” and “High BFP” gates in non-targeting and gene-targeting populations. Median of measurements from 3 replicates plotted. Boxes represent the median, IQR, and whiskers (±1.5 IQR). Two-sided pairwise t-tests with Bonferroni correction, **Padj ≤ 0.01, ****Padj ≤ 0.0001.

We first performed dCas9 ChIP-qPCR to confirm dCas9 binding at the targeted site in the GT CARGO, but not in the control non-targeting (NT) CARGO (Extended Data Fig. 6a). Our assay was designed to create an obstacle for elongating Pol II, and thus in our GT CARGO expressing cells, Pol II should accumulate downstream from the TSS and upstream from the targeted site. Indeed, Pol II ChIP-qPCR analysis showed increased Pol II levels upstream of the targeted site (Amplicons B and C), but not at the TSS (Amplicon A) (Fig. 6b). Interestingly, cohesin also accumulated at this site along with Pol II, as revealed by the RAD21 ChIP-qPCR (Fig. 6b). In contrast, in neutral control cell lines, we observed enrichment of dCas9 but not of Pol II or RAD21 signal in GT CARGO cells, indicating that dCas9 binding in the absence of transcription does not stall cohesin (Extended Data Fig. 6b).

Next, we assayed the consequences of these perturbations on gene expression. Consistent with the transcriptional block, in a large subpopulation of cells with GT CARGO, we observed a strong reduction of mTagBFP2 signal (“LowBFP”) (Fig. 6c, Extended Data Fig. 6c) alongside a “HighBFP” subpopulation, likely corresponding to cells with incomplete targeting of the locus (Fig. 6c, Extended Data Fig. 6c). We took advantage of this heterogeneity and gated the two subpopulations to analyze their eGFP signals. The LowBFP population had higher eGFP signal than HighBFP population, and this effect was more striking in the reverse orientation, which is associated with higher competition (Fig. 6d, Extended Data Fig. 6d). These results indicate that blocking transcription alleviates promoter competition and rescues endogenous *Sox2* eGFP levels, even in the presence of increased Pol II and cohesin at the integration site, demonstrating that cohesin accumulation is not sufficient to drive competition.

### Transcriptional read-through increases promoter competition

The observations that: (i) blocking transcription downstream of the TSS rescues competition despite Pol II pileup, and (ii) competition is dependent on promoter strength (and thus RNA output), suggested that the transcript itself may be driving the competition. We therefore asked whether transcript length also contributes. To promote transcriptional read-through, we removed the SV40 polyA signal from the reporter (Ef1a No-SV40pA) (Fig. 7a) and following integration, assayed alongside neutral and Ef1a (Ef1a wSV40 PolyA) reporters containing SV40 polyA (Fig. 7a). We observed decreased levels of mTagBFP2 protein in the No-SV40pA reporter cells (Fig. 7b). Since loss of the polyadenylation signal may affect mRNA stability or translation^44^, we also quantified the nascent *mTagBFP2* RNA levels by RT-qPCR. The polyA deletion had no effect on nascent *mTagBFP2* RNA level when the reporter was integrated in the reverse orientation, and caused a modest decrease in the forward integration (Fig. 7c). Loss of the polyA sequence from the reporter further decreased eGFP protein and mRNA levels (Fig. 7d, Extended Data Fig. 7c), consistent with increased competition.

**Figure 7.**
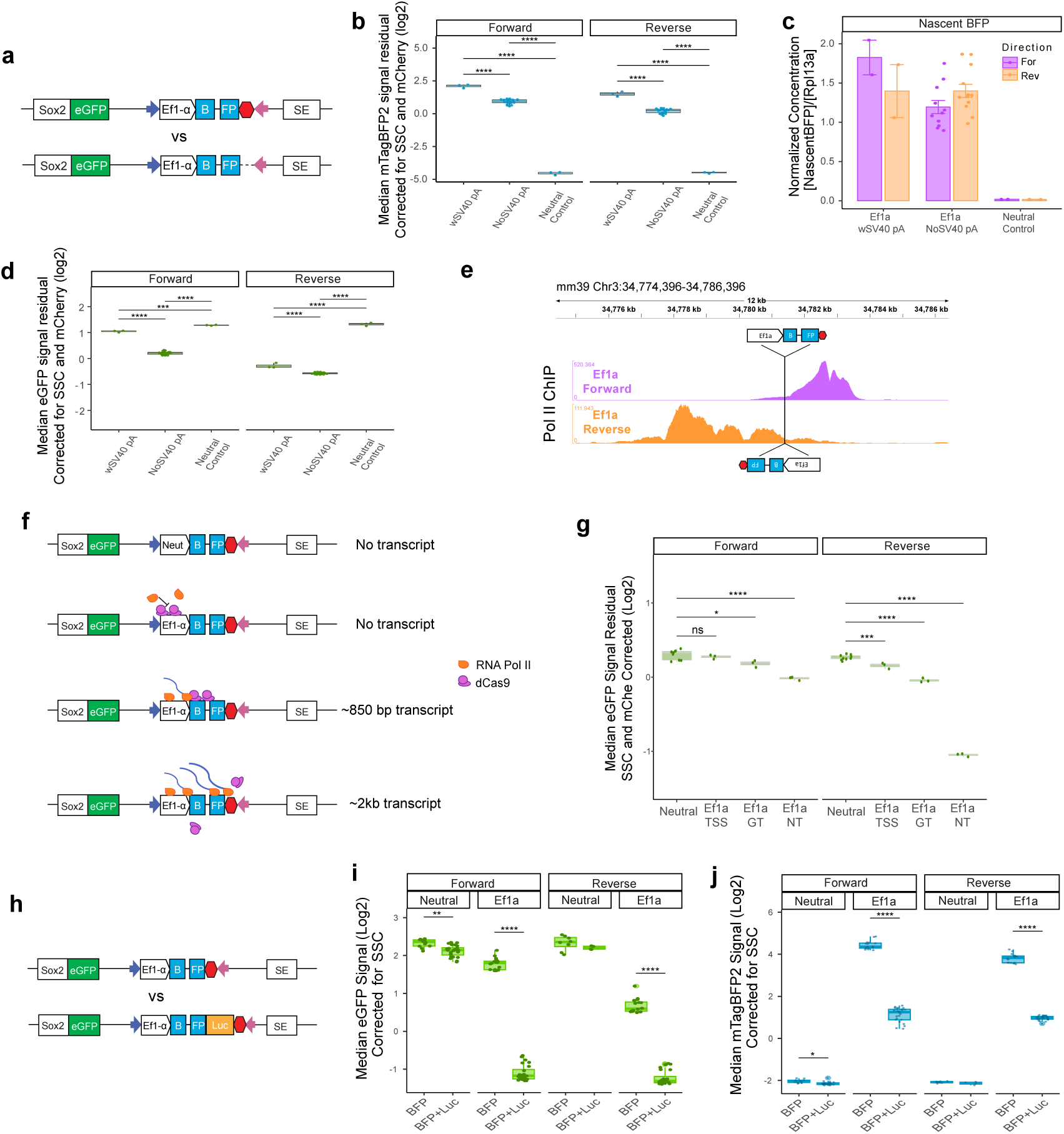
Transcript length affects competition. **a)** Schematic of Ef1a reporter integration with or without SV40 polyA signal (indicated in red). **b)** Corrected median mTagBFP2 signal from cells with Ef1a integrated with or without SV40 polyA. Two clonal lines with polyA were compared to 6 clonal lines without SV40 polyA. Flow measurements were repeated 3 times. Boxes represent the median, IQR, and whiskers (±1.5 IQR). Two-sided pairwise t-tests with Bonferroni correction, ****Padj ≤ 0.0001. **c)** RT-PCR analysis of nascent BFP transcript levels. Absolute concentration of Nascent BFP transcripts was normalized to housekeeping Rpl13a; 6 clones without SV40 polyA compared to 1 clone with Ef1a polyA and neutral polyA, in duplicate. mean ± s.e is plotted. **d)** Corrected median eGFP signal from cells with Ef1a reporter integrated with or without SV40 polyA. Two clones with polyA compared to 6 clones without SV40 polyA. Flow repeated 3 times. Boxes represent the median, IQR, and whiskers (±1.5 IQR). Two-sided pairwise t-tests with Bonferroni correction, ***Padj ≤ 0.001; ****Padj ≤ 0.0001. **e)** Total (non-allele specific) RNA Pol II ChIP-seq tracks zoomed near the landing pad in cells with Ef1a integrations. **f)** Schematic of dCas9-CARGO experiments for modifying transcript length. **g)** Corrected median eGFP signal from dCas9 CARGO experiment populations and Neutral controls. Experiment is repeated 3 times. Boxes represent the median, IQR, and whiskers (±1.5 IQR). One-sided pairwise t-tests with Bonferroni correction, *Padj ≤ 0.05; ***Padj ≤ 0.001; ****Padj ≤ 0.0001. TSS = Transcription Start Site Targeting dCas9-CARGO, GT = Gene-Targeting, NT=Non-Targeting. **h)** Schematic of Ef1a reporter integration with or without the addition of firefly luciferase coding sequence. **I,j)** Corrected median eGFP**(l)** and mTagBFP2**(j)** signal from promoter integration cells with BFP+Luciferase (BFP+Luc) compared to only BFP. 6 clones with BFP+Luc compared to 3 clones with only BFP. Flow repeated 3 times. Boxes represent the median, IQR, and whiskers (±1.5 IQR). Two-sided pairwise t-tests with Bonferroni correction, *Padj ≤ 0.05; ****Padj ≤ 0.0001.

We noted that the polyA removal caused a greater reduction of the eGFP signal in the forward direction (Fig. 7d, Extended Data Fig. 7c). We asked whether the difference in the transcriptional read-through length after the SV40 polyA signal could explain this observation. To test this, we performed RT-qPCR with primers tiling up to 10 kb downstream of the SV40 polyA site. In the forward orientation, removal of the polyA resulted in elevated transcript levels up to 5 kb downstream from the integrated reporter (Extended Data Fig. 7a). Importantly, we did not detect transcription 10 kb downstream, suggesting that transcriptional interference due to a head-on collision with the Pol II transcribing from the SE located 30 kb away is an unlikely explanation for the observed increase in competition (Extended Data Fig. 7a).

Surprisingly, in the reverse orientation, we detected transcription up to 5 kb downstream even in the polyA-containing reporter, with no further increases upon polyA deletion (Extended Data Fig. 7b). This is consistent with the smaller effect of polyA deletion in this orientation and suggests inherently longer read-through transcription in the reverse configuration. In agreement, Pol II ChIP-seq tracks showed orientation dependent differences in downstream Pol II distribution, with a more extended termination zone in the reverse orientation (Fig. 7e). These observations can explain why reverse insertions had lower mTagBFP2 levels (due to increased read-through and inverse correlation of transcript length and expression^45^) but yielded more competition than those in the forward direction. Indeed, deletion of the polyA signal in the Forward Ef1a reporter – which results in read-through of up to 5kb – attenuates eGFP signal to a degree comparable with the reverse Ef1a reporter containing the intact polyA (Fig. 7d, Extended Data Fig. 7c).

### Transcript length modulates levels of promoter competition

Next, we took two orthogonal approaches to directly test the association between competition and transcript length: (i) shortening the transcript via dCas9-CARGO, and (ii) lengthening the transcript by extending the coding sequence downstream of mTagBFP2.

First, we conducted a variant of our dCas9-CARGO experiment that allowed us to capture shortened transcript outputs (Fig. 7f). In a previous experiment, targeting dCas9 to the intron of the *mTagBFP2* gene (GT CARGO) (Fig. 6a) resulted in reduced competition (Fig. 6d). Given the GT CARGO position in the intron, this perturbation is predicted give rise to a short transcript of ∼ 850 bp. To eliminate transcription altogether, we designed another CARGO array, targeting the dCas9 molecules to the TSS of Ef1a. This led to a further rescue of GFP expression (almost to the level of neutral sequence insertion) (Fig. 7g). Comparing the effects of no transcript (in TSS CARGO), ∼850bp transcript (in GT CARGO) and ∼2.5kb transcript (in NT CARGO) on GFP levels, we observe that as the transcript length is reduced, the competition weakens (Fig 7g).

In a complementary approach, we lengthened transcripts in Ef1a and neutral test cassettes by adding a luciferase sequence downstream of mTagBFP2 (mTagBFP2–T2A–Luc), increasing the coding region from ∼1.3 kb to ∼3 kb (Fig. 7h), without including the readthrough. As expected, – given that at comparable initiation rates from the Ef1a promoter, less full-length transcripts are predicted to be produced per unit of time, – lengthening the coding sequence led to a reduction in the mTagBFP2 levels (Fig7h, Extended Data Fig. 7e). Despite this, we consistently observed dramatic reductions in the eGFP protein and RNA levels to a degree comparable with the SE deletion, whereas a neutral integration with similar length had modest effects on eGFP (Fig 7i, Extended Data Fig. 7d). Thus, transcript length is a major determinant of promoter competition.

### Patterns of promoter positioning across the genome in relation to strength

Having established determinants of promoter competition at a synthetic locus, we asked whether similar principles could have impacted genome organization and evolution. We reasoned that highly expressed promoters should preferentially reside near TAD boundaries, where they can contribute to insulation without interfering with long range enhancer-promoter interactions within the same regulatory domain. In agreement with prior observations that housekeeping genes, which are often highly expressed, are enriched at TAD boundaries^33^, we found that the likelihood of promoters being positioned near the boundary increased with expression (p<10^-16^) (Fig. 8a). Next, we modeled the location of promoter within the TAD as a function of its expression. We observed that the probability of the promoter to be located near either 5’ or 3’ boundary increased with its expression, with the most highly expressed genes more likely to be positioned at the boundary (Fig. 8b). While these observations do not disentangle correlation from causation, they are consistent both with the previous observations and with our competition model being generalizable and impacting evolution of regulatory domains across the genome.

**Figure 8.**
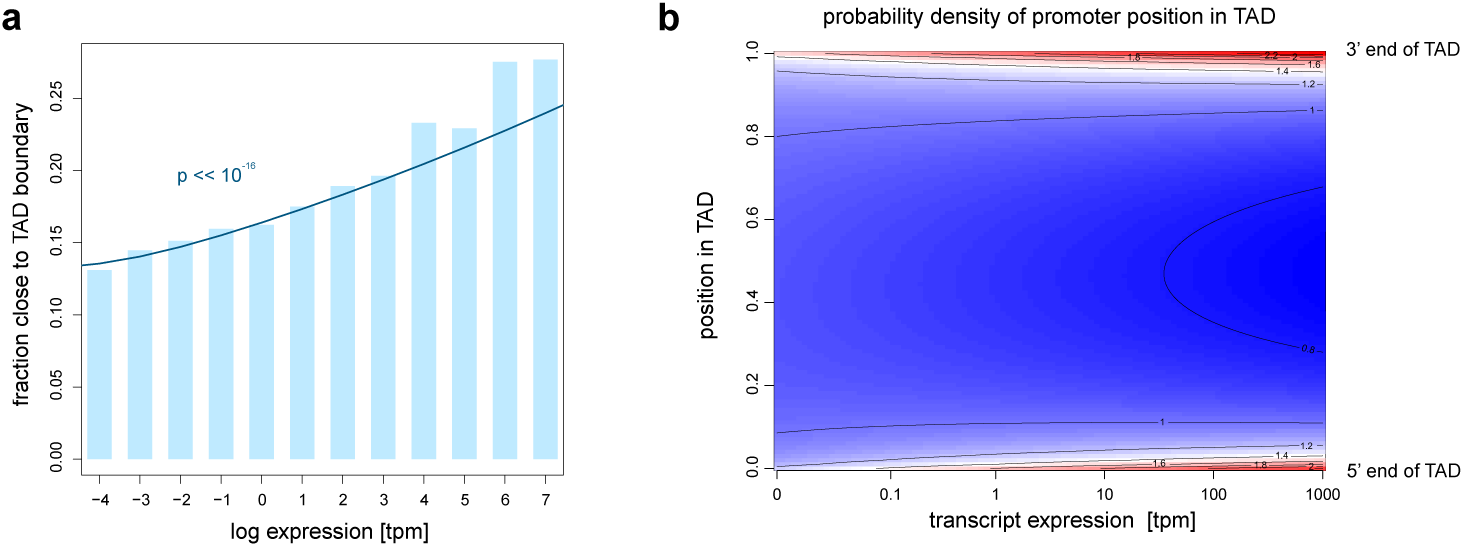
Probability of promoter positioning near a boundary increases with expression levels. **a)** Bar plot showing higher expressed genes are more likely to have promoters positioned closer to the TAD boundary. Promoter position within TAD was normalized to TAD length. Promoters positioned within (0.05-0.95) interval were considered as TAD-boundary distal, remaining as TAD-boundary-proximal. Plotted as blue rectangles is fraction of TAD-boundary proximal promoters for each bin of gene expression in mESCs. Navy line represents logistic regression line, where position within (0.05-0.95) interval was treated as failure modeled with log expression [tpm] as predictor variable. Statistical significance of the predictor was assessed using a two-sided Rao score test. **b)** Heatmap and contour plot of probability density function of beta distribution in x-axis for each gene expression value in y-axis. Promoter position was modeled with beta-regression with log expression [tpm] as predictor variable.

## DISCUSSION

In this work, we revealed promoter position, strength, and the associated transcriptional output as key determinants of promoter competition. Our findings argue against a simple enhancer resource-sharing model^46^ and instead support a mechanism in which transcription from a competing promoter can disrupt enhancer-promoter communication by promoting insulation. Our model predicts that such competition would arise when the competing promoter is intrinsically strong and positioned between the other promoter and its major enhancer. Beyond our system, studies where one promoter affects the output of another are consistent with this model. Additionally, promoters^47,48^ or simple transcribing units^49^ can serve an enhancer-blocking function, suggesting that transcription-driven insulation is a broader principle of genome organization. Nonetheless, transcription-dependent competition may also reflect broader effects on the regulatory hub structure^50,51^, and in such framework, the competition between promoters could be mutual and not strictly position-dependent.

While the effects of promoter competition measured in our system are modest, such dosage changes can have significant biological consequences, consistent with the widespread dosage sensitivity of haploinsufficient and triplosensitive genes^52,53^. Moreover, given that lengthening the transcript markedly increases competition, and endogenous mammalian transcripts are typically longer than even our extended reporters, our measurements may underestimate the competition potential. Thus, at endogenous loci, cis-regulatory activity must be calibrated to accommodate promoter competition or alternatively, promoter competition must be counteracted. Both are likely true, and in support of the latter, we observed HUSH-mediated silencing of the ectopic promoters. HUSH acts co-transcriptionally and targets transcribed non-self DNA, such as retroviruses or transposable elements^20,23,25,54^, potentially providing a first line of defense against deleterious promoter competition arising from de novo retroelement promoter insertions. However, this does not explain how competition among endogenous promoters within the same regulatory domain is resolved. Our data show that promoter strength and position govern competition, suggesting that placing the highly active promoters away from the centers of regulatory domains may minimize regulatory conflicts. Indeed, TAD boundaries have long been known to be enriched for promoters of housekeeping genes^33^ or active retroelements^54^. More recently, developmental genes have also been shown to be enriched near the TAD boundaries and contribute to insulation^47^. Our results indicate that rather than housekeeping-developmental distinctions, it is the promoter strength and the level and length of its associated transcript that determine insulating and competitive capacity. In this context, the generally short and unstable nature of many noncoding transcripts^55^ may serve to limit this competitive potential.

Our study shows that inserting a transcribing unit driven by a strong promoter is sufficient to induce insulation and thus promoter competition. Importantly, this effect does not arise from the insert itself, the associated Pol II machinery, or its potential to stall cohesin. Rather, it requires transcriptional output, with transcript level and length as key determinants of competition strength. Although the underlying mechanism remains unclear, one potential explanation is RNA-mediated regulation of transcriptional condensates ^56,57^. Superenhancers, such as the one studied here, nucleate transcriptional condensates enriched in transcription factors and cofactors^58,59^. It has been posited that while low levels of RNA may support condensate formation, higher levels of longer RNA, associated with elongation can dissolve them^56,57^. This provides a natural explanation for the length dependence in competition, as longer RNAs may more effectively disrupt multivalent interactions and dissolve condensates, by displacing IDR-rich coactivators. Consistent with this model, increased competition is associated with reduced Sox2–SE contact frequency in Micro-C, suggesting that insulation and condensate disruption may reflect the same underlying process at different resolutions. A second, non-mutually exclusive explanation of our findings is transcription-coupled chromatin remodeling. Transcription can partially unwrap nucleosomal DNA, evict histones, reposition nucleosomes and generate supercoiling, any of which could conceivably alter CRE interactions. Elongation through longer transcription units would extend these effects over larger regions, potentially changing the biophysical properties of the intervening chromatin fiber and reducing enhancer-promoter contact probability. Both proposed mechanisms are consistent with the dependence of competition on transcript length and output.

Altogether, the more abundant and longer a competing promoter transcript, the greater its interference with endogenous gene expression. While future studies will be needed to dissect the contributions of RNA, chromatin remodeling or other potential underlying mechanisms, our results establish parameters governing promoter competition within complex regulatory domains.

## Acknowledgements

We thank B. Ren and H. Huang (UCSD) for sharing the Sox2 landing pad cell lines used in this study; M. Fuller, A. Kundaje and members of the Wysocka lab for feedback on the project; S. Tabatabaee, R. Fueyo, C. Jensen, T. Chern, S. Bar, M. Dunjic, E. Magee, K. Brennan, O. Crocker, L. Ichino and C. Ni for critical reading of the manuscript. This work was supported by HHMI, R35 GM131757 and Lorry Lokey endowed professorship (to J.W.), R01EB035127, R01CA300848 (to A.S.H.), NSF EF2022182 (to A.B.), Daiichi Sankyo Foundation of Life Science Fellowship and HFSP Long-Term Fellowship (LT0022/2024-L) (to M.N.), and Gerhard Casper Stanford Graduate Fellowship (to M.K.).

## Author Contribution Statement

M.K. and J.W. conceived the project. M.K. performed all experiments and analysis except for Region Capture Micro-C, which was performed and analyzed by M.N. T.S. performed genome wide analysis on promoter positioning. J.W., A.B., A.S.H. supervised the project. M.K. and J.W. wrote the manuscript with feedback from all authors.

## Competing Interest Statement

Authors declare no competing interests.

## METHODS

### Cell Culture

F123 mESCs were a gift from Bing Ren’s laboratory at UCSD^21^. Once received, the cell lines were further validated for presence of the eGFP and mCherry fluorophore tags and landing pad locations with genotyping. Fluorophore signal was also validated by flow cytometry. Chromosomal abnormalities were ruled out by comparing coverage from ChIP inputs to input from known proper diploid cells. The cells were cultured as previously described^60^. In brief, cells were grown in monolayer on tissue culture plates pretreated with 7.5 ug/ml poly-L-ornithine (Sigma, P4638) in PBS and 5 ug/ml laminin (Gibco, 23017015) in PBS consecutively for at least 1 hour at 37°C. Cells were maintained in serum-free 2i + LIF media (500mL DMEM/F-12 (Gibco, 11320082), 1x Gem21 Neuroplex w/o Vitamin A (Gemini Bio, 400-161-010), 1x N2 Neuroplex (Gemini Bio, 400-163-005), 2.5g BSA (Gemini Bio, 700-104P), 1xMEM NEAA (Thermo, 11140050), 1xSodium pyruvate (Thermo, 11360070), and 1xAnti-Anti (Sigma, A5955) containing MEK inhibitor PD0325907 (0.8 uM, Selleck, S1036), GSK3b inhibitor CHIR99021 (3.3 mM, Selleck, S2924), and leukemia inhibitory factor (LIF, produced in-house). Media was changed every day, cells were passaged every 2-3 days when 80-90% confluent using 0.25% Trypsin-EDTA (Gibco, 25200072) to release cells from the plate. Dissociation reaction was quenched with quenching media (500mL DMEM/F-12, 1x Anti-Anti(Sigma), 10% FBS (Gemini Bio,100-106)). Cells were re-plated on PLO/Laminin plates at 1:6-1:20 dilutions.

R1 mESCs was used as unstained control, or to generate compensation controls expressing a single fluorophore, and were also cultured in 2i+LIF media. The wild type cell line was provided by Andras Nagy’s lab at Lunenfeld-Tanenbaum Research Institute, Canada and authenticated by karyotyping. All cell lines were routinely tested for mycoplasma.

### Donor plasmid cloning for RMCE

The donor vector was adapted from pGL3-Basic plasmid (Promega, E1751). First, pGL3 was modified to remove luciferase and SV40 polyA signal. Two ultramers containing FRT and FRT3 (IDT), and a gene-fragment with a multiple cloning site (MCS), followed by mTagBFP2 with a mouse b-globin intron and an SV40 polyA site (Twist Biosciences) were ordered with matching restriction sites. The empty pGL3 backbone was re-ligated with these three components to create parental donor plasmids. To allow for inserts in different orientations, two versions of the donor plasmid were made with the same strategy (for pDonor-For, FRT-mTagBFP2-FRT3, for pDonor-Rev, FRT3-mTagBFP2-FRT). 1kb fragment of promoters were amplified from mouse genomic DNA with nested PCR using Platinum SuperFi II polymerase (Thermo, 12361010) and neutral fragment was amplified directly from DH5-alpha cells (NEB, C2987H) with overhangs matching the MCS. Ef1a (Addgene #55632) and CAG (Addgene #161974) were taken from published plasmids. For Ef1a allelic series and Rpl13a mutant experiments, the fragments were ordered from Twist. For noSV40 PolyA experiments parental donor plasmids were modified to remove the SV40 polyA with a 20bp neutral fragment, followed by Ef1a promoter insertion into the MCS. For assembling donor plasmids with luciferase coding sequence, donor plasmids with only mTagBFP2 were digested, oligos with T2a sequence were annealed to yield appropriate restriction site overhangs and luciferase CDS was amplified from pGL3 plasmid with restriction site overhangs. All plasmids were verified with analytical digest, followed by whole plasmid sequencing (Plasmidsaurus). All promoter sequences are listed in Supplementary Table 1 and all primers used in Supplementary Table 3.

### Landing pad – RMCE cell lines

Parental landing pad lines (LP_bw or LP_ds) were co-transfected with a Flippase expression plasmid (Addgene #89574) and donor plasmid with promoter at 1:2 molar ratio using Lipofectamine 2000 (Invitrogen, 11668019) according to manufacturer protocol. Cells were allowed to recover for 7-10 days after transfection and passaged onto 10cm plates with media containing 0.5uM ganciclovir (Sigma, SML2346). Surviving isolated clones were manually picked and transferred to a 48-well plate. After another week in culture, cells were passaged for maintenance and genomic DNA collection with Quick Extract (Lucigen, QE09050). Clones were genotyped with primers specific to forward or reverse inserts using OneTaq Master Mix with GC buffer (NEB, M0489L). At least three independent clones were kept for each promoter insert for subsequent flow cytometry analysis.

### Generation of HUSH Knockout Cell lines

To stably express targeting guides and Cas9, px459(Addgene, 62988) plasmid was cloned into a PiggyBac backbone to generate pb459. Two guides targeting MPP8 and two non-targeting^25^ control guides were cloned into pb459 using golden gate assembly. One clone from each promoter insert was selected and transfected with either two Mpp8 targeting or two non-targeting guide plasmids, alongside Supertransposase (SBI, PB210PA-1) in 1:1:1 equimolar ratio using lipofectamine 2000 according to manufacturer recommendations. Next day populations were passaged onto 2i+LIF media containing 1ug/ml puromycin (Invivogen, ant-pr-1). The parental clones were grown alongside the Mpp8 deletion and non-targeting controls and assayed together in flow cytometry experiments.

To establish clonal MPP8 deletion lines, the same two MPP8 targeting guides were cloned into px459. Ef1a and neutral insert lines were transfected with px459 plasmids. Next day populations were passaged onto 2i+LIF media containing 1ug/ml puromycin. After 3 days of selection, 1000-2000 surviving cells were plated on a 10cm plate coated with 5ug/ml fibronectin (Sigma, FC01010MG) in PBS. After 1 week of culture, individual clones were picked, genotyped. Selected clones were validated with western blot for MPP8 knockout. All guides and primers used for genotyping are listed in Supplementary Table 3.

### Generation of SE deletion lines

To delete the SE in an allele specific manner, two guides with PAM sequences overlapping CAST SNPs were chosen. Two SE targeting guides were cloned into px459 and Ef1a insert lines were transfected with these plasmids. Next day populations were passaged onto 2i+LIF media containing 1ug/ml puromycin. After 3 days of selection, surviving cells were sorted in Sony MA900 cell sorter (housed in Stanford Stem Cell FACS Facility) for loss of eGFP and maintenance of mCherry signal. 1000-2000 sorted cells were plated on a 10cm plate coated with 5ug/ml fibronectin in PBS. After 1 week of culture, individual clones were picked and later genotyped. At least three independent clones were kept for each SE deletion for subsequent flow cytometry analysis. All guides and primers used for genotyping are listed in Supplementary Table 3.

### Generation of SE CTCF motif mutation lines

To mutate the putative CTCF motifs in the SE in an allele specific manner, two guides with PAM sequences overlapping CAST SNPs were chosen. Two SE targeting guides were cloned into px459. Next, three DNA fragments (800-1.2kb in length) of a portion of the SE with the CTCF motifs mutated, with appropriate restriction enzyme sites were ordered from Twist Biosciences. The fragments were cloned into a pGL3 backbone with a selection counterselection (LoxP-Ef1a-HyTK-LoxP) insertion. Cell lines with Ef1a insertion either in the in-between or downstream landing pad were transfected with this donor plasmid and two px459 constructs with the appropriate guides. Next day populations were passaged onto 2i+LIF media containing 150ug/ml hygromycin (Invivogen, ant-hg-1). After 5 days of selection, the cells were sorted to confirm there was no effect on the mCherry allele. 1000-2000 sorted cells were plated on a 10cm plate coated with 5ug/ml fibronectin in PBS, still in hygromycin containing media. After 1 week of culture, individual clones were picked and genotyped with primers from the inside of the selection cassette to outside the homology arms to confirm mutation of the CTCF motifs. Successful clones were transfected with a pCAG-CreO-Puro plasmid to excise the selection cassette and selected with 1ug/ml puromycin for two days after transfection. After expansion, the clones were genotyped once again to confirm the removal of the selection/counterselection cassette and put in ganciclovir containing media to kill any cells that might still have the cassette. Next, clones were transfected with MPP8 targeting guides as described before to create population of HUSH-KO cells. At least two independent clones were kept for each promoter insert for subsequent flow cytometry analysis. SE sequence containing mutated CTCF sites is listed in Supplementary Table 1. All guides and primers used for genotyping are listed in Supplementary Table 3.

### Generation of Rad21-dTag lines

A guide^61^ targeting the C-terminal of Rad21 was cloned into px459. A donor plasmid with ∼1.1kb Rad21 homology arms flanking FKBP12^F36V^-Flag-T2a-Puro was generated with Gibson assembly. Ef1a insertion lines were co-transfected with donor and px459 plasmid at a 1:1 molar ratio. Next day populations were passaged onto 2i+LIF media containing 1ug/ml puromycin. After 3 days of selection, 1000-2000 surviving cells were plated on a 10cm plate coated with 5ug/ml fibronectin in PBS and kept in puromycin containing media. After 1 week of culture, individual clones were picked and genotyped. Successful clones were confirmed with DMSO vs 500nM dTag treatment for 6 hrs followed by western blot to assure maintenance of tagged Rad21 levels and proper degradation. All guides and primers used for genotyping are listed in Supplementary Table 3.

### CARGO Array and Cell Population Generation

CARGO 6-mers were generated as described previously^42,62^. Briefly, CARGO constant region was digested from pGEMT-hU6-SL. Custom oligos were annealed and phosphorylated. Each annealed oligo independently was combined with the digested CARGO constant region to generate minicircles. Minicircles were then treated with Plasmid-safe exonuclease to remove unligated fragments according to the manufacturer’s protocol. Minicircles were then combined and cleaned using Zymo DNA Clean & Concentrator-5 (Zymo, D4014) kit. Separately CARGO plasmid backbone was generated from a modified version of pK335 (Addgene, #227505) plasmid was generated with no florescence reporter, pK335-NoColor with restriction digest. CARGO array was then generated by combining the minicircles and the CARGO plasmid backbone at a 3:1 molar ratio in a golden gate reaction. Assembly reaction was then treated with Plasmid-safe exonuclease per the manufacturer’s protocol. 1uL of this reaction was used to transform 25uL NEB Stable competent E.coli (NEB, C3040H). Successful preps were confirmed with full plasmid sequencing (Plasmidsaurus).

Clonal Ef1a or neutral insert lines were transfected with PhiC Integrase (SBI, FC200PA-1 and either TSS targeting, mTagBFP2 Gene targeting or non-targeting CARGO in equimolar ratio. Next day, cells were passaged onto 2i+LIF media containing 200ug/ml G418 (Invivogen, ant-gn-1). After 7 days, populations were transfected with pB-dCas9-Puro plasmid. Next day, cells were passaged onto 2i+LIF media containing 200ug/ml G418 and 1ug/ml puromycin. A no-dCas9 expression control was also performed but all comparisons are between Targeting and Non-targeting populations with dCas9 expression. Cells were then grown and expanded for 10 days in double selection media. On 11^th^ day, a portion of cells were crosslinked for ChIP and the remainder were processed with flow cytometry. Independent populations were generated as described each time for 3 biological replicates. All guide RNA sequences and custom oligos for CARGO synthesis are provided in Supplementary Table 3.

### Flow cytometry data acquisition and analysis

For promoter screens such as ones depicted in Fig1E, Fig 2D, Supp 2B, Supp 3B, cells were grown on 48-well plates. Cells were detached from plates with trypsin and resuspended with 2i+LIF media containing 5% serum (Gemini, 100-525). Resuspended cells were mixed thoroughly and transferred to a U-bottom 96 well plate. Then, cells were analyzed in a LSR Fortessa analyzer (BD) housed in Stanford Stem Cell FACS facility, in High Throughput Sampler mode. eGFP, mCherry and mTagBFP2 signal were recorded for at least 10000 cells for each clone, with hardware compensation. For experiments with less than 30-40 samples, cells grown in 6 or 12-well plates were released with trypsin, quenched and spun down. The pellet was resuspended with cold FACS buffer (1xPBS, 5% FBS), cells were strained through 0.35uM mesh for subsequent analyses in LSR Fortessa, in tube mode and at least 50000 events were collected.

The FCS files were processed in FlowJo (v10.10.0) and gated for singlets (consecutively gated for SSC.A vs FSC.A, FSC.H vs FSC.A, FSC.W vs FSC.A, SSC.H vs SSC.A, SSC.W vs SSC.A). The linear values from gated populations were exported as csv files and further analyzed in R (v4.3.3). The files were imported into R for downstream analysis.

Flow cytometry measures integrated fluorescence per cell, and in this system the eGFP/mTagBFP2 signal is influenced by cell size and by variation in Sox2 promoter activity. To account for these confounders, we regressed fluorescence measurements on log-transformed side scatter (SSC), which correlates with cell size, and on log-transformed mCherry levels, which report on insertion-independent Sox2 promoter activity and also correlate with cell size. The regression was performed across all single-cell measurements for each experiment using the model: lm(log2(Flurophore) ∼ log2(SSC)+ log2(mCherry). For SE deletion experiments, only SSC was included in the regression because wild type cells with no fluorophores were used as control comparison. Residuals from these models were used for all downstream analyses and visualization.

If a promoter insertion clone had reduced mCherry levels (signifying differentiation) or bimodal eGFP signal (duplication of the insert allele or mixed clones) measured by flow cytometry, despite passing genotyping quality controls, the clone was omitted from the analysis and replaced with a new clone when possible.

### Luciferase reporter assays

Ef1a mutant promoters were cloned into pGL3-Basic (Promega, E1751) using restriction digestion. Two distinct preps for each mutant promoter were cleaned with Plasmid-Safe ATP-Dependent DNase (Lucigen, E3101K). Transfections were performed in 24-well plates, with each well receiving 10ng of pGL3 plasmid, 0.5ng of control pRL firefly renilla plasmid (Promega E2261), 89.5 mL carrier DNA (circularized pGEMT plasmid) and 0.25ul of Lipofectamine in 50ul Optimem (Gibco 31985070). 24 hours after transfection, cells were assayed with Dual-Luciferase Reporter Assay System (Promega, E1960) according to manufacturer’s recommendation. Briefly, cells were washed in PBS, and lysed in 100 uL 1X passive lysis buffer (in PBS) for 10 min. 20 uL lysate was then transferred to an opaque flat-bottomed plate for reading with a luminometer (Promega, GloMax). The automated injector added 100 uL LARII reagent and the well was read using the following parameters: 2 s delay, 10 s integration. 100 uL Stop-and-Glow reagent was then injected into the well and read using the same parameters. Luciferase assays were repeated twice, using biological duplicates (two distinct preps of same promoter vector), transfected in duplicate; empty vector and SV40 promoter were included in each experiment as controls.

### Total cell lysis and Western Blot

Cells were washed once with PBS before lysing on-plate in ice (1 mL per 10 cm plate) with ice cold RIPA buffer (50 mM Tris-HCl pH 7.4, 150 mM NaCl, 1 mM NP40, 0.5% sodium deoxycholate, and 0.1% SDS) with 1x cOmplete EDTA-free protease inhibitor cocktail (Roche, 11873580001) and 1mM PMSF. The lysate was centrifuged for 10 minutes at max speed (>15,000 x g) at 4°C. The supernatant was moved to a new tube and protein concentration was quantified using the Pierce BCA protein kit (Thermo Scientific, 23227) and normalized. Samples were then denatured by addition of 1x NuPAGE LDS Sample Buffer (Invitrogen, NP0007) and 1.25% b-mercaptoethanol and subsequently heating to 70° C for 10 min. Samples were loaded to 4-20% Novex Tris-glycine gels (Invitrogen) and run at 100V for 30min, then 180V for 40min in Tris-glycine buffer (25 mM Tris and 192 mM glycine) with 0.1% SDS. Gels were transferred onto nitrocellulose membranes (GE Healthcare, 10600003) for 1 h at 400 mA in Tris-glycine buffer with 20% methanol. The membrane was stained with 0.1% Ponceau S in 3% trichloroacetic acid, then blocked with 5% milk and 1% BSA in PBS with 0.1% Tween-20 (PBST) for 30 min at room temperature and then incubated with primary antibody overnight at 4°C followed by horseradish peroxidase (HRP)-conjugated secondary antibody incubation for 1 h at room temperature, with 4 washes of PBST after each antibody incubation. Chemiluminescence was performed with Amersham ECL Prime reagent (Cytiva, RPN2232) and imaged with an Amersham ImageQuant 800 (Amersham). Membranes were then incubated for 1 h at RT with HRP-conjugated loading control antibodies, washed and imaged again. Antibodies used include Mpp8 (Proteintech, 16796-1-AP), Rad21 (Abcam, ab992), b-actin-HRP (Abcam, ab49900), Hsp90-HRP (Cell Signaling, 79641S), Goat-Rabbit IgG (H+L) HRP (ProteinTech, SA00001-2).

### ChIP-qPCR & ChIP-Seq

10-30 million sub-confluent cells were grown on 6 cm (for H3K9me3, CTCF) or 10 cm (for Pol II, Rad21, dCas9) plates, washed with 1x PBS and fixed with freshly prepared 1% methanol-free formaldehyde in PBS for 10 minutes at room temperature with nutation. Glycine was added to a final concentration of 125 mM to quench for an additional 10 minutes with nutation. The quenched formaldehyde solution was removed, and cross-linked cells were scraped from plate in ice cold PBS with 0.001% Triton-X100. Cells were pelleted at 1350xg for 5 minutes at 4°C, washed with ice cold PBS, and pelleted as before. All liquid was removed, pellets were snap-frozen in liquid nitrogen and stored at -80°C until further processing.

On the day of the immunoprecipitation, pellets were thawed on ice for 30 minutes and resuspended in 5 mL LB1 (50 mM HEPES-KOH pH 7.5, 140 mM NaCl, 1 mM EDTA, 10% glycerol, 0.5% NP-40, 0.25% Triton X-100, with 1X Roche cOmplete Protease Inhibitor Cocktail and 1mM PMSF), rotated end over end at 4°C for 5 minutes, then pelleted at 1350xg for 5 minutes at 4°C. The pellet then was resuspended in 5mL LB2 (10 mM Tris-HCl pH 8.0, 200 mM NaCl, 1 mM EDTA, 0.5 mM EGTA, with 1X Roche cOmplete Protease Inhibitor Cocktail and 1mM PMSF) and again rotated end over end at 4°C for 5 minutes, then pelleted at 1350xg for 5 minutes at 4°C. Lysates were then resuspended in ice cold LB3 (10mM Tris-HCl, pH 8.0, 100mM NaCl, 1mM EDTA, 0.5 EGTA, 0.1% Na-Deoxycholate, 0.5% N-laurylsarcosine with 1x Roche cOmplete Protease Inhibitor Cocktail and 1mM PMSF) and transferred to appropriate tubes and incubated on ice for 10 minutes. For samples processed for ChIP-seq, 1ml LB3 was used, and sample was transferred to Covaris AFA tubes and sonicated for 10 minutes at peak power = 140, duty factor = 10, and cycles per burst = 20 in the E220 evolution Covaris. For samples that were being processed for ChIP-qPCR 0.3ml LB3 was used, and sample was transferred to Diagenode TPX tubes. ChIP-qPCR samples were sonicated for 8 cycles of 30s on/ 30s off on high power using Bioruptor Plus (Diagenode), additional cycles added until chromatin peak was observed around 500bp. To quantify total chromatin amount, and determine whether additional cycles of sonication were necessary, 5 ul out of 300ul (or 5 out of 1ml) were taken out and the rest of the sample kept on ice. To each 5uL sample, 195uL elution buffer (1% SDS, 0.1M NaHCO3), 2ul 5M NaCl, and 1ul RNaseA (0.2mg/ml final) was added. Samples were incubated at 65°C for 1hour with shaking. 1ul ProteinaseK (0.2mg/ml final) was added and samples were incubated at 65°C for an additional 1hour with shaking. DNA was then extracted using Zymo ChIP Clean & Concentrator-5 (Zymo, D5205) kit according to the manufacturer’s protocol. Portion of eluted DNA was run on an agarose gel to see sheared chromatin size and if desired peak at 500bp was achieved, DNA was quantified with Qubit DS High Sensitivity Kit (Thermo, Q33231). If chromatin peak was above 500 bp, additional cycles of sonication performed and then validation and quantification step repeated. The remaining sonicated lysate was diluted to 1mL in LB3 and spun at max speed (>15000xg) for 10 minutes at 4°C. Supernatant was transferred to a fresh DNA LoBind tube and Triton-X100 was added to a final concentration of 1%. Based on Qubit measurements, chromatin was normalized and 5% input samples were reserved and stored at -20°C. For H3K9me3 10ug, for Rad21 ChIP-seq samples 20ug, for Pol II ChIP-seq samples 15ug, for CTCF, Rad21, Pol II and Cas9 ChIP-qPCR samples 20ug chromatin was used. For each IP, 5ug antibody was added to 1ml soluble chromatin sample. Then samples were rotated vertically overnight (12-16hr) at 4°C. Antibodies used include H3k9me3(Abcam, ab8898), Rad21 (Abcam, ab992), RNA Pol 2 (Biolegend, 664906), Cas9 (Active Motif, 61757), CTCF (Abcam, ab128873).

The next day, 100uL Protein G Dynabeads (Thermo, 10004D) per ChIP sample were washed 3 times with ice cold blocking buffer (0.5% BSA(w/v)(Sigma, A3294) in PBS) then ChIPs were added to beads and incubated with vertical rotation for 4-6 hours at 4°C. ChIPs were then washed 5 times with RIPA buffer(50mM HEPES-KOH, pH 7.5, 500mM LiCl, 1mM EDTA, 1% NP-40, 0.7% Na-Deoxycholate), once with TE + 50mM NaCL (50mM Tris-HCl, pH 8.0, 10mM EDTA, 50mM NaCl), then eluted from the bead in 200uL elution buffer(1% SDS, 0.1M NaHCO3) at 65°C for 30min and eluate was transferred to a new DNA LoBind tube. To reverse cross-link and degrade RNA, 8uL 5mM NaCl and 2uL RNaseA (0.2mg/ml final) was added to eluate and to thawed input samples that were diluted to 200uL in elution buffer. All samples and inputs were incubated overnight (12-16hr) at 65°C with shaking. ProteinaseK (0.2mg/ml final) was added to samples and inputs, which were incubated for another 2 hours at 55°C with shaking. For ChIP-seq, DNA was extracted with phenol chloroform extraction and for ChIP-qPCR samples were processed using Zymo Clean& Concentrator-5 kit. DNA was then stored at -20°C.

For ChIP-seq, libraries were prepared using the NEBNext Ultra II DNA kit (NEB, E7645S) using up to 50 ng of input or ChIP DNA, with ∼4-8 cycles of amplification, with no pre-PCR size selection but a post-PCR double-sided 0.5x/0.9x Ampure XP bead clean-up. Final quantification and QC were assessed by Qubit and Tapestation (Agilent), respectively, and libraries were pooled for sequencing on NovaSeq X Plus (Illumina) platform.

For ChIP-qPCR, DNA samples were diluted by a factor of 5. Master mixes containing 5uL/reaction Sensifast SYBR No-Rox 2x mix (Meridian Bioline, 98020), 0.25ul 10mM primer mix, and 0.75uL water were prepared and kept on ice. 6uL of appropriate qPCR master mix was added to each well of a 384 well plate followed by 4uL of diluted cDNA. All samples were run in technical triplicate in a LightCycler 480 (Roche).

### ChIP-seq Analysis

An N-masked genome and a SNP file for Cast_EiJ and 129/Sv were generated using SNP annotation VCF file from the Mouse Genomes Project (Release V8, 12/2021) with the SNPSplit^63^ (v0.6.0) in dual strain mode. The reads were trimmed using skewer^64^ (v0.2.2) and aligned to the N-masked mouse genome using bowtie2 ^65^ (v2.5.1). After alignment the reads were filtered for quality score and PCR duplicates were removed. Then, the reads were assigned to either Cast or 129 allele with SNPSplit. For visualization, RPGC normalized allele specific bigwig files were generated using deeptools ^66^ (v3.5.1) bamCoverage.

### Analysis of Public ChIP-seq data for promoter features heatmap

Public datasets (SRAs listed in Supplementary Table 2) were downloaded with sratoolkit (v3.0.7). Adaptors were trimmed using Fastp^67^ (v0.21.0) and Fastq files were aligned with bowtie2 (v2.5.1) to mm39. The reads were filtered for quality score and PCR duplicates were removed from only paired-end datasets. Bam files were normalized and converted to bigwig files with bamCoverage from deeptools (v3.5.1) using RPGC normalization. All bigwig files were then analyzed together with multiBigwigSummary from deeptools to create a coverage matrix over 1kb regions of endogenous promoters. The coverage values were log transformed. Scaled values were plotted as a heatmap of relative enrichment.

### RT-qPCR

Cells were grown in 12-well plates. For Rad21-dTag experiments, cells were treated with 500nM of dTag^v^-1 or DMSO for 6 hours. For each experiment, more than 5 biological replicates (clones) were collected and assayed twice. For BFP+Luc lengthening experiments, 3 biological replicates from each insert were used. Prior to collection, cells were washed with PBS at RT, then 400uL Trizol reagent (Invitrogen, 15596026) was added to each well. Plates were nutated for 5 minutes at RT and samples were then transferred to RNase free tubes and stored at -80°C until further processing.

To extract RNA, samples in Trizol were thawed on ice, then total RNA was extracted using Zymo Direct-zol (Zymo, R2052) kit and cDNA was prepared from 2ug RNA using High Capacity cDNA Reverse Transcription Kit (Invitrogen, 4368813) according to manufacturer’s protocol. cDNA samples were diluted by a factor of 20 prior to RT-qPCR. Master mixes containing 5uL/reaction Sensifast SYBR No-Rox 2x mix (Meridian Bioline, 98020), 0.25ul 10mM primer mix, and 0.75uL water were prepared and kept on ice. 6uL of appropriate qPCR master mix was added to each well of a 384 well plate, followed by 4uL of diluted cDNA. All samples were run in technical triplicate using a LightCycler 480 (Roche).

### Analysis of qPCR data

All qPCR primers were validated prior to use in all experiments with the requirement to generate a single PCR product as determined by melt curve analysis and to have an efficiency of 1.8-2.0. Standard curves for all validated primer sets were obtained and used to convert Cp values into relative concentrations. Primers are provided in Supplementary Table 3.

All individual qPCR replicates are the average of technical triplicates. All ChIP-qPCR samples are quantified as percent input recovery. For RT-qPCR sample concentrations were normalized to a housekeeping gene, Rpl13a, then normalized by expression for a control line neutral insert for transcript length experiments that was normalized by Rpl13a. For Rad21 dTag experiments, Gapdh was used as a housekeeping control, and housekeeping normalized samples were then normalized by mCherry expression in their respective DMSO control counterpart. For experiments in Fig7c and Supp7, sample concentrations were only normalized to Rpl13a and not normalized further to a cell line, to display absolute levels of transcript, not relative levels.

### Region capture Micro-C (RCMC) library generation

RCMC libraries were prepared as previously described with minor modifications^27,32^. For each biological replicate, 15 million cells per sample were processed as 3 tubes of 5 million cells each. Cells were collected by trypsinization and crosslinked with 3 mM disuccinimidyl glutarate (DSG; Thermo Fisher, 20593) for 35 min at room temperature with gentle rotation, followed by addition of 1% formaldehyde (Thermo Fisher, 28908) for 10 min. Reactions were quenched with Tris-HCl at pH7.5(Sigma-Aldrich, T1503) to a final 0.375 M, and crosslinked cells were pelleted at 950 × g for 7 min and washed in PBS supplemented with 1% BSA (Thermo Scientific, 15260037).

Nuclei were resuspended in ice-cold Micro-C buffer (MB)#1 (50 mM NaCl, 10 mM Tris-HCl pH 7.5, 5 mM MgCl2, 1 mM CaCl2, 0.2% NP-40 alternative (Millipore, 492018), 1× protease inhibitor cocktail (Sigma, 5056489001)) for 20 min and centrifuged at 1,750 × g for 5 min. After one wash with the same buffer, chromatin was digested with 25 U of micrococcal nuclease (MNase; Worthington, LS004798) at 37 °C for 20 min, quenched with EGTA to a final 4 mM, and heat-inactivated at 65 °C for 10 min. Samples were then washed twice in MB#2 (50 mM NaCl, 10 mM Tris-HCl pH 7.5, 10 mM MgCl_2_, 100 µg/mL BSA (Sigma, B8667)).

Digested nuclei were phosphorylated with T4 polynucleotide kinase (NEB, M0201) in NEBuffer 2.1 with 2 mM ATP and 5 mM DTT at 37 °C for 15 min. Klenow fragment (NEB, M0210) was added to a concentration of 0.5 U/µL at 37 °C for 15 min to generate blunted ends, followed by filling-in with biotin-dATP and biotin-dCTP (Jena Bioscience, NU-835-BIO14 and NU-809-BIOX) together with dGTP and dTTP (Jena Bioscience, NU1003 and NU1004) at 66 µM each. After incubation at 25 °C for 45 min, reactions were quenched with 30 mM EDTA, followed by enzyme inactivation at 65 °C for 20 min. Nuclei were pelleted at 2,000 × g for 5 min, washed once with MB#3 (50 mM Tris-HCl pH 7.5, 10 mM MgCl2). Ligation was performed in 1× T4 DNA ligase buffer (100 µg/mL BSA) with T4 DNA ligase (NEB, M0202; 10,000 U per tube) at 25 °C overnight. Residual biotin on unligated ends was removed with exonuclease III at 37 °C for 15 min (NEB, M0206), followed by reverse crosslinking overnight at 65 °C in 1% SDS and 250 mM NaCl with proteinase K (final 2 mg/mL; Viagen Biotech, 501-PK) and RNase A (final 0.1 mg/mL; Thermo Fisher, EN0531).

DNA was purified using Zymo DNA Clean & Concentrator-25 (Zymo, D4034), separated on a 1% agarose gel (120 V, 60 min), and the 200–400 bp dinucleosome band was excised and purified (Zymo, D4008). Biotinylated ligation products were captured on T1 streptavidin beads (Invitrogen, 65601). From bead-bound DNA fragments, Micro-C libraries were generated using NEBNext Ultra II DNA Library Prep (NEB, E7645S) on-bead following the manufacturer’s instructions, and amplified by 8 cycles of PCR followed by 0.95x AMPure XP bead purification (Beckman Coulter, A63881).

Custom 80-mer, biotinylated oligonucleotide panels (Twist Bioscience) were designed to cover the Sox2 locus (chr3:33,750,000-35,650,000 in mm39 coordinates), the inserted promoter sequences, as well as the eGFP and mCherry transgenes. For capture, the Micro-C libraries above corresponding to each kind of promoter insertion were pooled by replicate, and 2–4 µg total library was used per capture. Hybridization followed the Twist Standard Target Enrichment v2 protocol (https://www.twistbioscience.com/resources/protocol/twist-target-enrichment-standard-hybridization-v2-protocol) using Universal Blockers (Twist, 100578), Mouse Cot-1 DNA (Invitrogen, 18440016), and Hybridization Mix (Twist, 10587). The hybridized products were captured with streptavidin beads (Twist, 100983), washed, and PCR-amplified using Equinox Library Amplification Mix (Twist, 104126) for 12 cycles. Following 1.8x AMPure XP bead purification (Beckman Coulter, A63881), the libraries were pooled and sequenced on a single lane of NovaSeq X Plus 25B (paired-end 2 × 150).

### RCMC data processing

RCMC data were processed largely as described previously with minor adjustments for our capture panels^68^. All custom scripts are available on GitHub (see Code Availability).

Briefly, all analyses were carried out against sample-specific hybrid references that contain both 129S1 and CAST haplotypes together with the engineered insertions. Homozygous SNPs for 129S1 and CAST were applied to the mm39 reference to generate per-haplotype FASTA files, appending _129 and _CAST to sequence headers, which were then concatenated to form a combined 129/CAST reference used for alignment and downstream phasing. These references were indexed with bwa v0.7.19-r1273 for alignment^69^. Paired-end FASTQ files were aligned to the corresponding custom references with bwa-mem using the -SP option. The SAM outputs were converted to BAM with samtools v1.21.

BAM alignments were converted to 2D contact pairs with pairtools v1.1.2 in a haplotype-resolved workflow^70^. First, we parsed read pairs (pairtools parse) with a minimum mapping quality of 0, annotated the mapq, XA, NM, AS, and XS fields, discarded SAM records, masked read walks, and supplied chromosome sizes, yielding *.unphased.XA.pairs.gz. We then phased pairs with pairtools phase using the XA tag (--tag-mode XA) and appended the allele-specific suffixes _129 and _CAST, producing *.phased.XA.pairs.gz. Phased pairs were coordinate-sorted with pairtools sort. Finally, duplicates were identified and marked with pairtools dedup (Cython backend; --max-mismatch 1), while retaining the haplotype annotations as extra pair-level columns (phase1, phase2). The resulting *.phased.sorted.XA.nodup.pairs.gz files were used for downstream analyses.

After the aforementioned phasing with pairtools phase (tag-mode XA with the suffix configuration mapping 0 to 129S1 and 1 to CAST) and deduplication, we used the per-side phase tags (phase1, phase2) to derive two allele-resolved subsets. First, a double-sided (DS) subset retained only contacts for which both ends were phased to the same allele—i.e., 129S1 if phase1 == 0 and phase2 == 0, or CAST if phase1 == 1 and phase2 == 1. Second, a single-sided (SS) subset was defined more permissively to include contacts where at least one end carried the allele-informative phase call while the mate could be unphased—i.e., for 129S1 we kept (0,0), (0,.) or (.,0); for CAST we kept (1,1), (1,.) or (.,1). Each subset was written out as .129_allele.{ds|ss}.pairs and .CAST_allele.{ds|ss}.pairs. To restrict analyses to the Sox2 capture interval on chr3, we retained only pairs with both ends inside the capture analysis window (mm39 chr3:33,750,000–35,653,652) and excluded pairs flagged as duplicates (pairtools pair_type == DD). Reads overlapping excluded intervals corresponding to the engineered insertion sites were also removed: mm39 chr3:34,705,521–34,706,292 (Sox2 fluorescent-protein insertion) and 34,782,089–34,785,059 (promoter insertion).

For each sample and replicate, we converted pair files into single-resolution .cool contact matrices with cooler v0.10.3, providing explicit column mappings and CAST-normalized chromosome sizes^71^. Specifically, we ran: cooler cload pairs --assembly mm39_129S1_CAST -c1 2 -p1 3 -c2 4 -p2 5 <custom.chrom.sizes>:1000 <PAIRS> <out.1000.cool>. This was performed for allele-specific DS and SS subsets. We then generated multi-resolution, balanced .mcool files with cooler zoomify --resolutions 1000,2000,5000,10000,20000,50000,100000,200000,500000 --balance -o <out.mcool> <in.1000.cool>. Biological replicates were merged at the .cool level using cooler merge, and the merged products were re-zoomified to create pooled .mcool files. Contact-map visualization from balanced .mcool files, loop-strength quantification (Sox2 promoter, inserted promoters, and the Sox2 control region [SCR]), and insulation-score calculation were performed with custom Python scripts. All analyses used 5 kb bins; insulation was computed with cooltools v0.5.4 using cooltools.api.insulation.calculate_insulation_score with a 100 kb window (--res-bin 5000, --win-bp 100000, ignore_diags=2, min_dist_bad_bin=2)^34^.

### Promoter Position Analysis

Promoter position within TAD was normalized to TAD length. Promoters positioned within (0.05-0.95) interval were considered as TAD-boundary distal, remaining as TAD-boundary-proximal. The fraction of TAD-boundary proximal promoters for each bin of gene expression in mouse ESC (defined as floor log (expression +.02)) were plotted as a barplot. Navy line represents logistic regression line, where position within (0.05-0.95) interval was treated as failure modeled with log (expression + 0.02) as predictor variable with lm(formula = close ∼ log(expression + 0.02), family = binomial()). Next, position of promoter was modeled with beta-regression with log (expression + 0.02) as predictor variable. A heatmap and contour plot of probability density function of beta distribution in abscissa for each gene expression value in ordinate were plotted. The actual model used is (position ∼ strand | log(expression + 0.02)

### Statistics and reproducibility

All statistical analyses were performed using R (v4.3.3). No statistical method was used to predetermine sample size. Sample size, number of replicates, statistical tests and definition of error bars were stated in the legends. No data were excluded from data analysis, except for flow measurements from differentiated clones as explained in detail above. The experiments were not randomized. The investigators were not blinded to allocation during experiments and outcome assessment.

## Data availability

Genomic datasets have been deposited at GEO and are publicly available as of the date of publication under GSE296229 (ChIP-Seq) and GSE312207 (RCMC).

Plasmids generated are deposited at Addgene (Plasmids# 253414- 253442).

Source data is available with this manuscript. Raw data and cell lines are available upon request from

## Code availability

Custom scripts used to analyze allele-specific RCMC are available at https://github.com/ahansenlab/Koska-et-al-RCMC_PromoterCompetition_analysis_code

---

**Extended Data Fig. 1.**
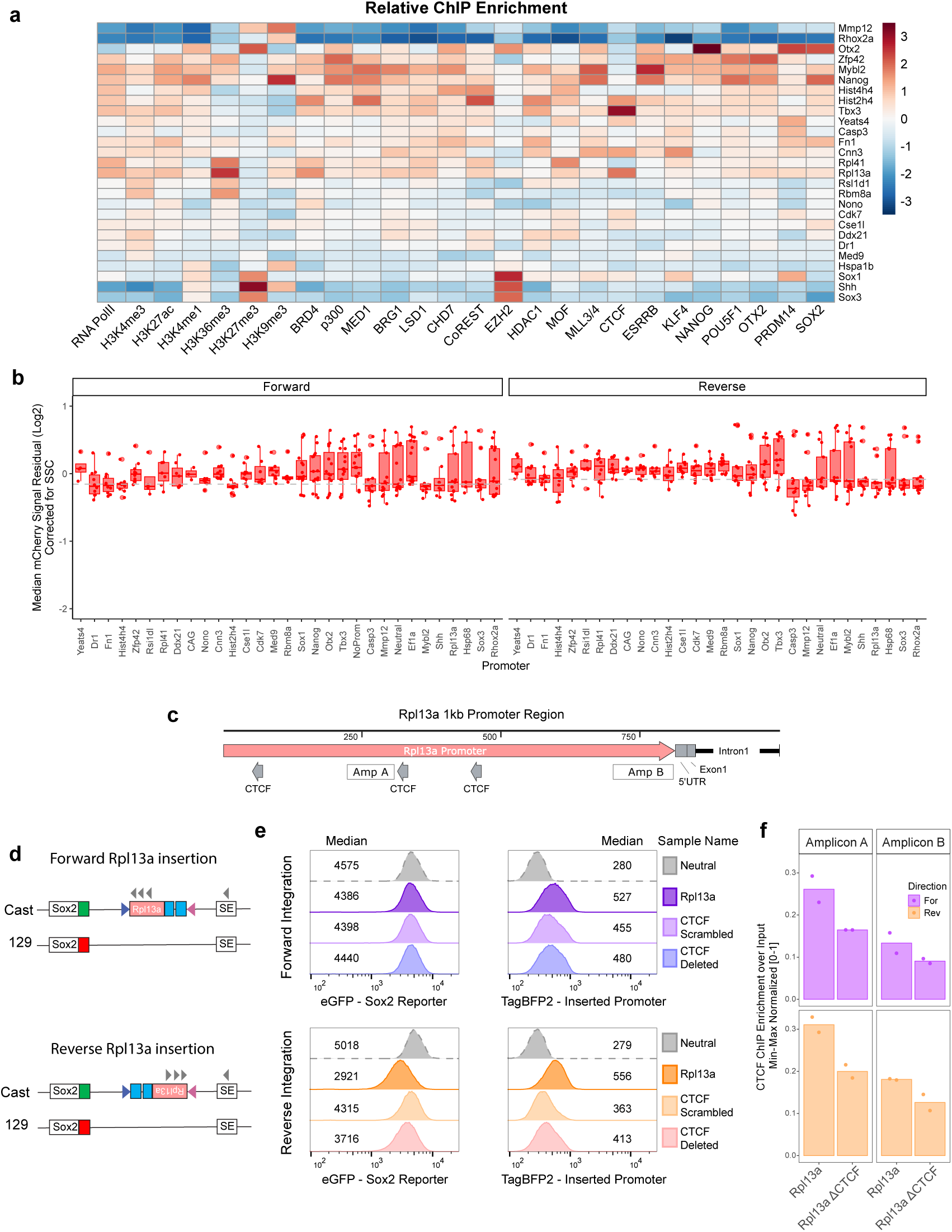
Additional information on selected promoters and screen outliers. **a)** Heatmap showing relative normalized ChIP-seq enrichment of histone modification, TF and COF binding profiles at 1 kb promoter regions used in the screen. Fill represents log coverage scaled by columns. **b)** Boxplot of median mCherry residual signal for each promoter after correction for SSC with points of each independent observation. Data includes 3 biological (from three independent clonal lines) and 3 technical replicates for each promoter. Boxes represent the median, IQR and whiskers (±1.5 IQR), outlier displayed as large red dots. **c)** Schematic of Rpl13a promoter region used in this study and CTCF motifs. Direction of the arrow shows orientation of the CTCF binding sites. White filled boxes represent amplicon regions for ChIP-qPCR. **d)** Schematic of Rpl13a promoter integrated in the landing pad. Arrows show directionality of CTCF sites in Rpl13a and SE. **e)** Flow cytometry measurement histograms of eGFP and mTagBFP2 for Rpl13a mutant promoters integrated at the landing pad. Numbers represent median measurements. **f)** CTCF ChIP-qPCR Min-Max normalized percent input recovery at sites labeled in (b) for WT and ΔCTCF Rpl13a insert lines. Note that signal from endogenous Rpl13a promoter alleles is also detected with this amplicon; n=2, bars represent means, individual measurements displayed as dots.

**Extended Data Fig. 2.**
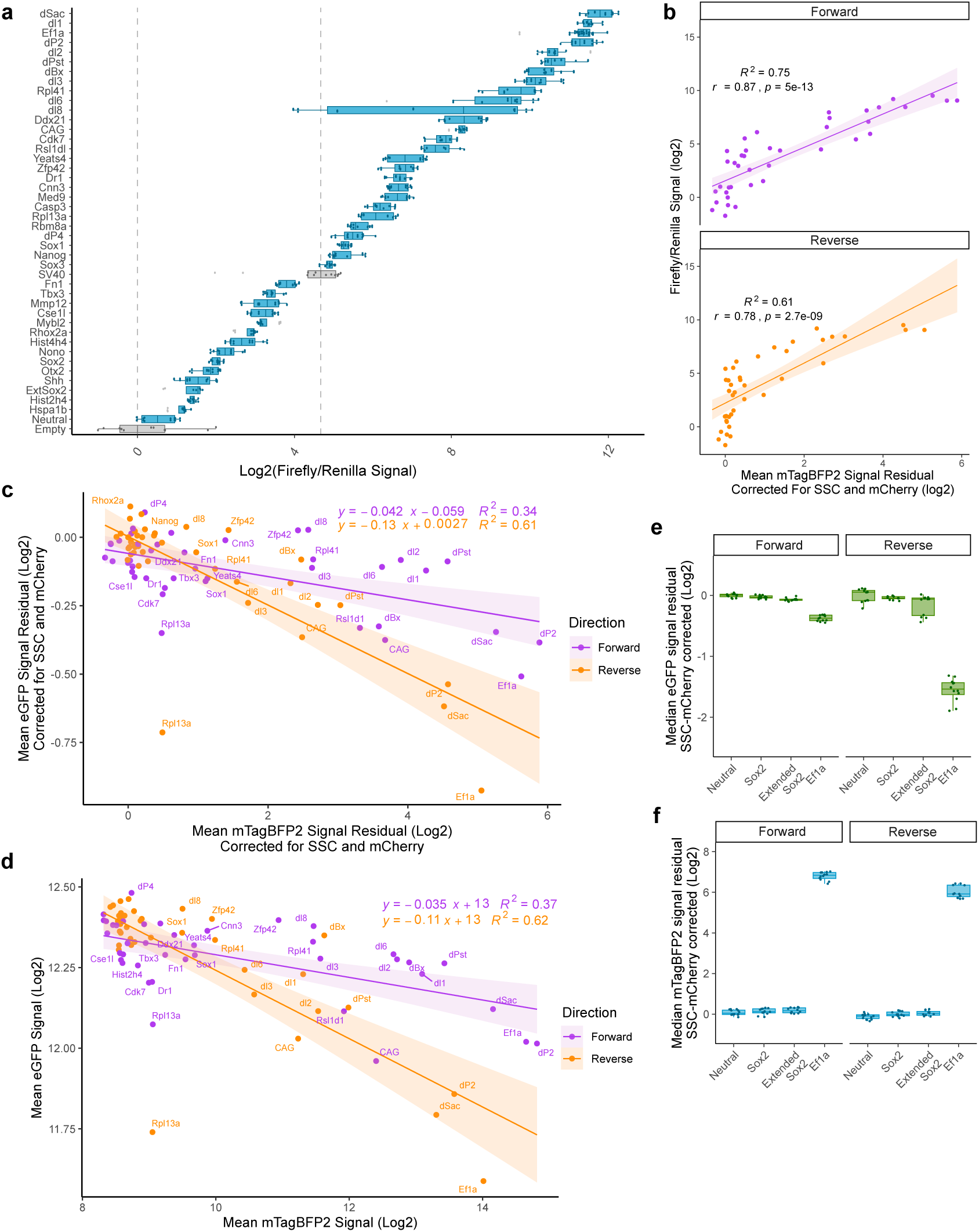
Promoter competition is driven by the strength of the competing promoter. **a)** Episomal Luciferase reporter assay measurements from each promoter. Measurements performed twice with two distinct preps of reporter plasmid, transfected in duplicate. Firefly divided by Renilla signal is displayed. Boxes represent the median, IQR, and whiskers (±1.5 IQR), outlier measurements displayed in gray. Boxplots for control measurements with empty vector and SV40 promoter are filled with gray. Dotted line represents the median of the SV40 and Ef1a measurements. **b)** Correlation plot of promoter strength measured in the episomal luciferase reporter assay compared to strength of the integrated promoter measured by mTagBFP2 output. Coefficient of determination R^2^ values for linear regression, Pearson’s correlation coefficient R and two-sided p value for correlation are displayed. **c)** Scatterplot of corrected mean eGFP and mTagBFP2 signal of all promoters tested, for 3 biological replicates (independent clonal lines), measured in triplicate. Equation and R^2^ value for linear regression for each direction is displayed. Values shifted vertically and horizontally to make neutral insert adjusted to 0. **d)** Same scatterplot as (c), with uncorrected raw mean eGFP and mTagBFP2 signal for all biological and technical replicates. **e, f)** Corrected median eGFP and mTagBFP2 measurements from lines with Sox2 promoter and controls. Median from 3 biological replicates (independent clonal lines), measured in triplicates plotted together. Boxes represent the median, IQR, and whiskers (±1.5 IQR).

**Extended Data Fig. 3.**
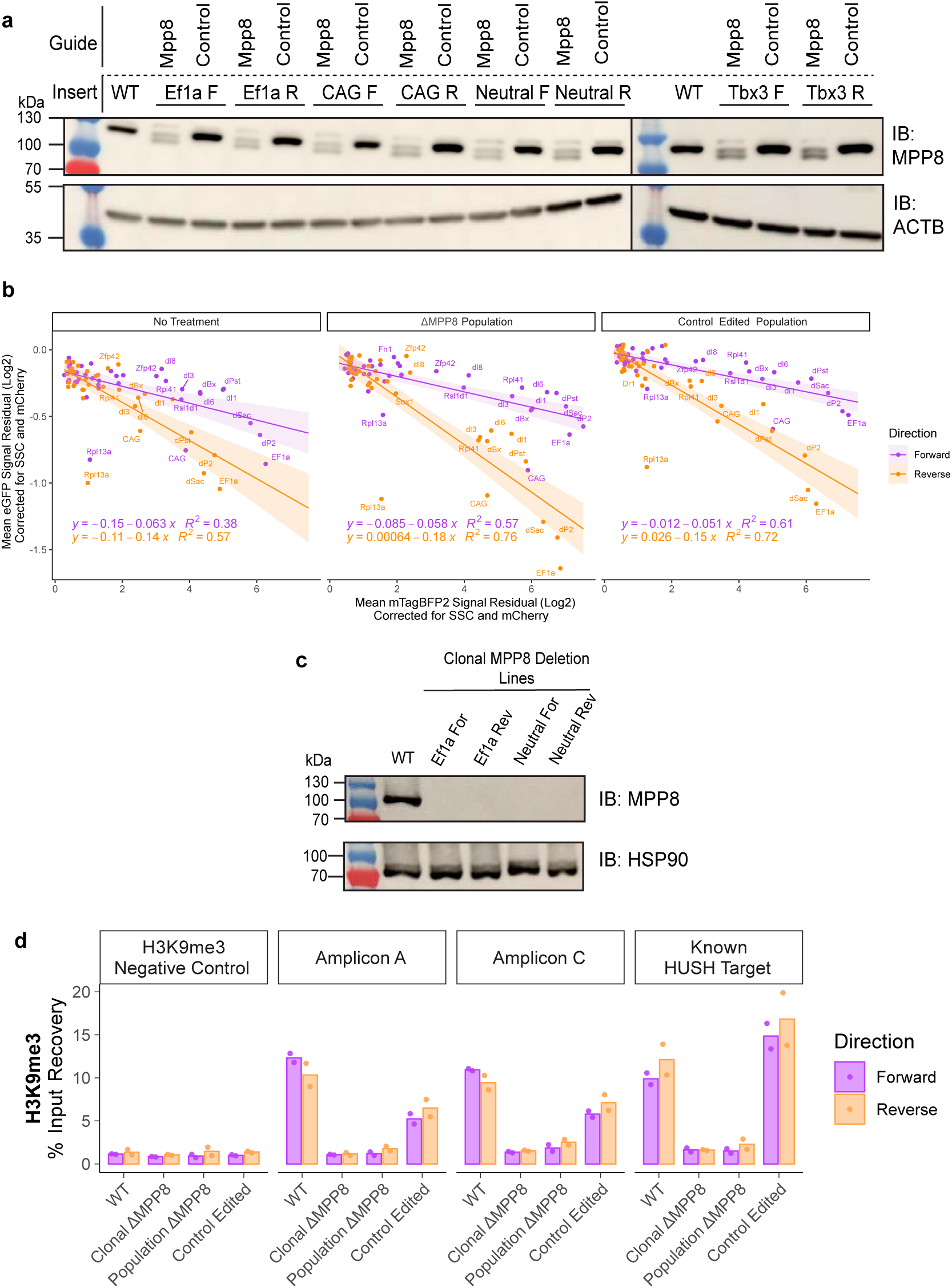
Controls for HUSH ablation in landing pad cells. **a)** MPP8 western blot in WT, and a subset of MPP8 targeted, and Control targeted populations. Membranes were probed with antibodies against MPP8 and ACTB as a loading control. Representative of two independent experiments. **b)** Scatterplot of corrected eGFP and mTagBFP2 signal for all promoters tested in WT, ΔMPP8 and control edited backgrounds. ΔMPP8 and control edited measurements were done in triplicates in population of cells. **c)** MPP8 western blot in WT, and clonal ΔMPP8 lines. Membranes were probed with antibodies against MPP8 and HSP90 as a loading control. Representative of two independent experiments. **d)** H3K9me3 ChIP-qPCR percent input recovery at sites labeled in (3B) for Ef1a insert lines in clonal WT, clonal ΔMPP8, population ΔMPP8 and control edited populations. n=2, bars represent means.

**Extended Data Fig. 4.**
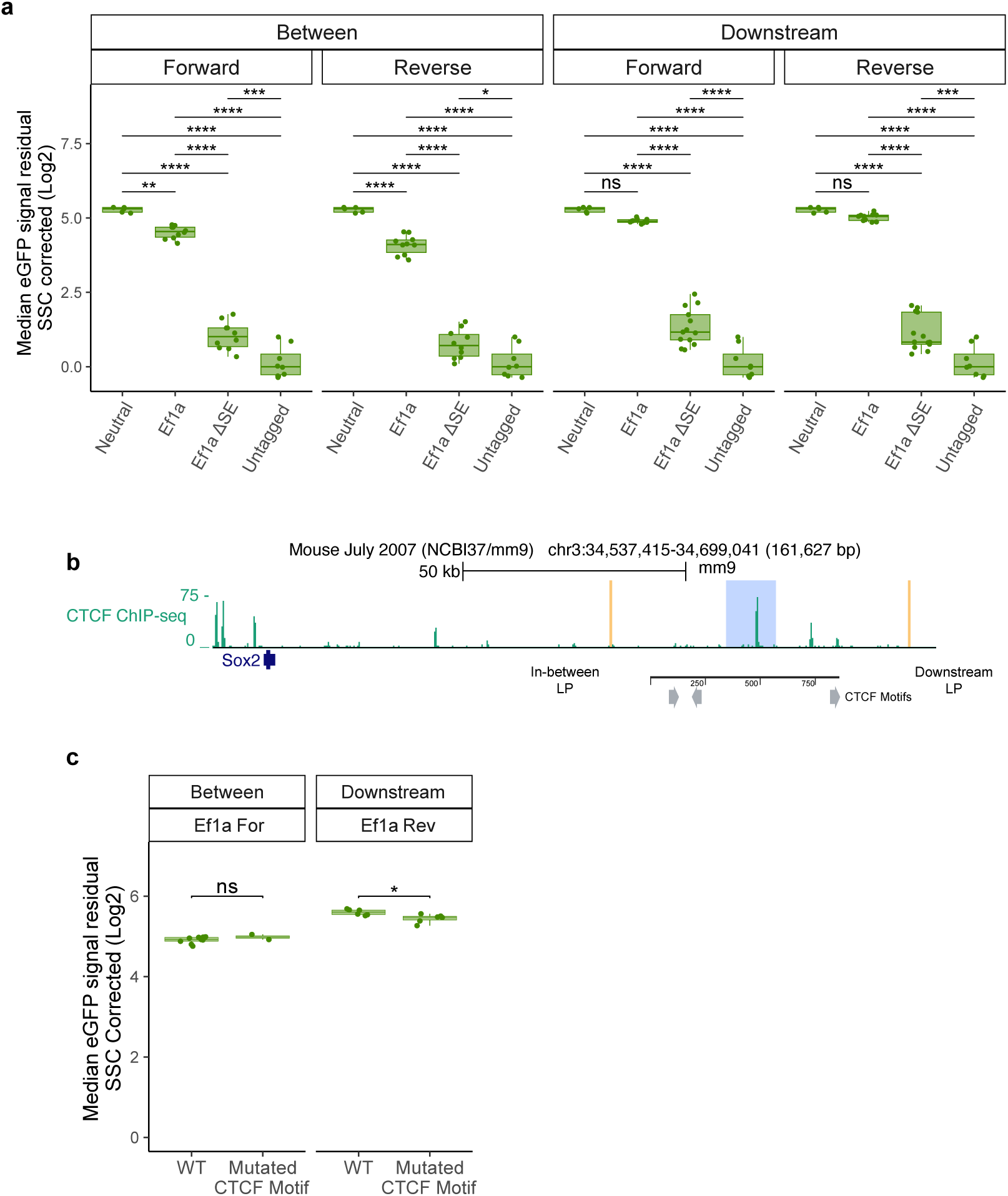
Changes in eGFP levels in response to SE deletion. **a)** Corrected median eGFP measurements from in between and downstream landing pad lines with neutral insert integrated, Ef1a integrated, Ef1a integrated with ΔSE and untagged WT control. Median from 3 biological replicates (independent clonal lines), measured in triplicates plotted together. Values shifted vertically to make untagged control adjusted to 0. Boxes represent the median, IQR, and whiskers (±1.5 IQR). Two-sided pairwise t-tests with Bonferroni correction, *Padj ≤ 0.05; **Padj ≤ 0.01; ***Padj ≤ 0.001; ****Padj ≤ 0.0001. **b)** ChIP-seq tracks for CTCF at Sox2 locus. Landing pad locations are highlighted in orange and SE in blue. Putative CTCF motifs underlying the most prominent CTCF peak shown in gray, direction of the arrow shows orientation of the CTCF binding sites. **c)** Corrected median eGFP measurements from cell lines with Ef1a reporter inserted in either in-between or downstream landing pad with CTCF motifs mutated in the SE. Median from 2 biological replicates (independent clonal lines), measured in triplicates plotted together. Boxes represent the median, IQR, and whiskers (±1.5 IQR). Two-sided pairwise t-tests with Bonferroni correction, *Padj ≤ 0.05.

**Extended Data Fig. 5.**
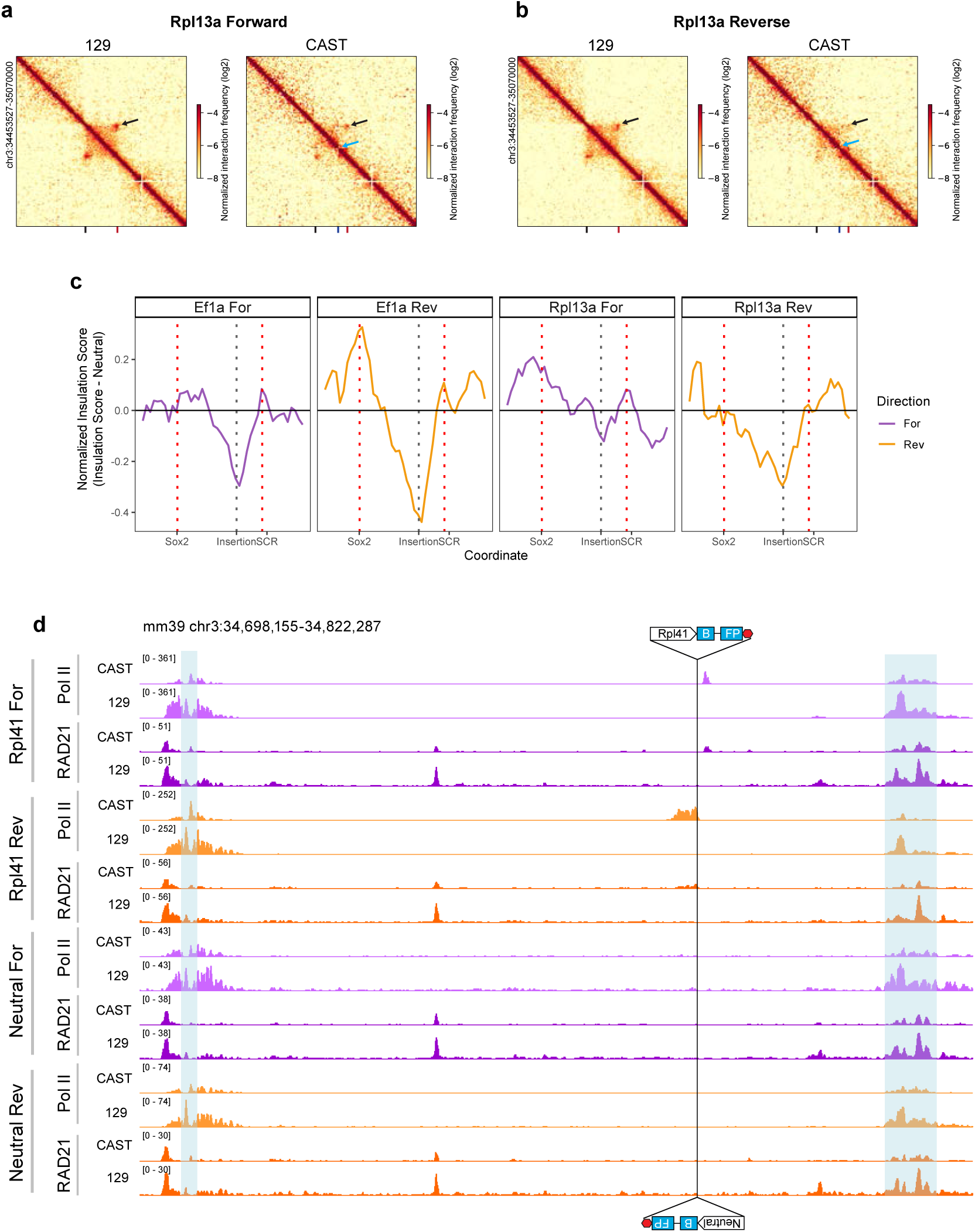
Cohesin accumulates near the Rpl41 but not the near neutral sequence insert. **a,b)** Contact map visualization of RCMC data for Rpl13a For (a) and Rpl13a Rev (b) integrations at 5-kb resolution. Contacts map for control allele 129 shown on the left, for insert allele CAST on the right. Sox2 location is marked with a black notch, insert with blue, and SE with red. Black arrows highlight SE-Sox2 contact, blue arrows highlight domain split. Maps generated with merged reads from two biological replicates (independent clones). **c)** Insulation scores for CAST alleles normalized to Neutral insert. **d)** Allele specific RNA Pol II and Rad21 ChIP-seq tracks in the Rpl41 and Neutral sequence integration cell lines. Landing pad location and direction marked with cartoon of Rpl41 and Neutral control integration, blue highlights mark Sox2 coding sequence and SE, respectively.

**Extended Data Fig. 6.**
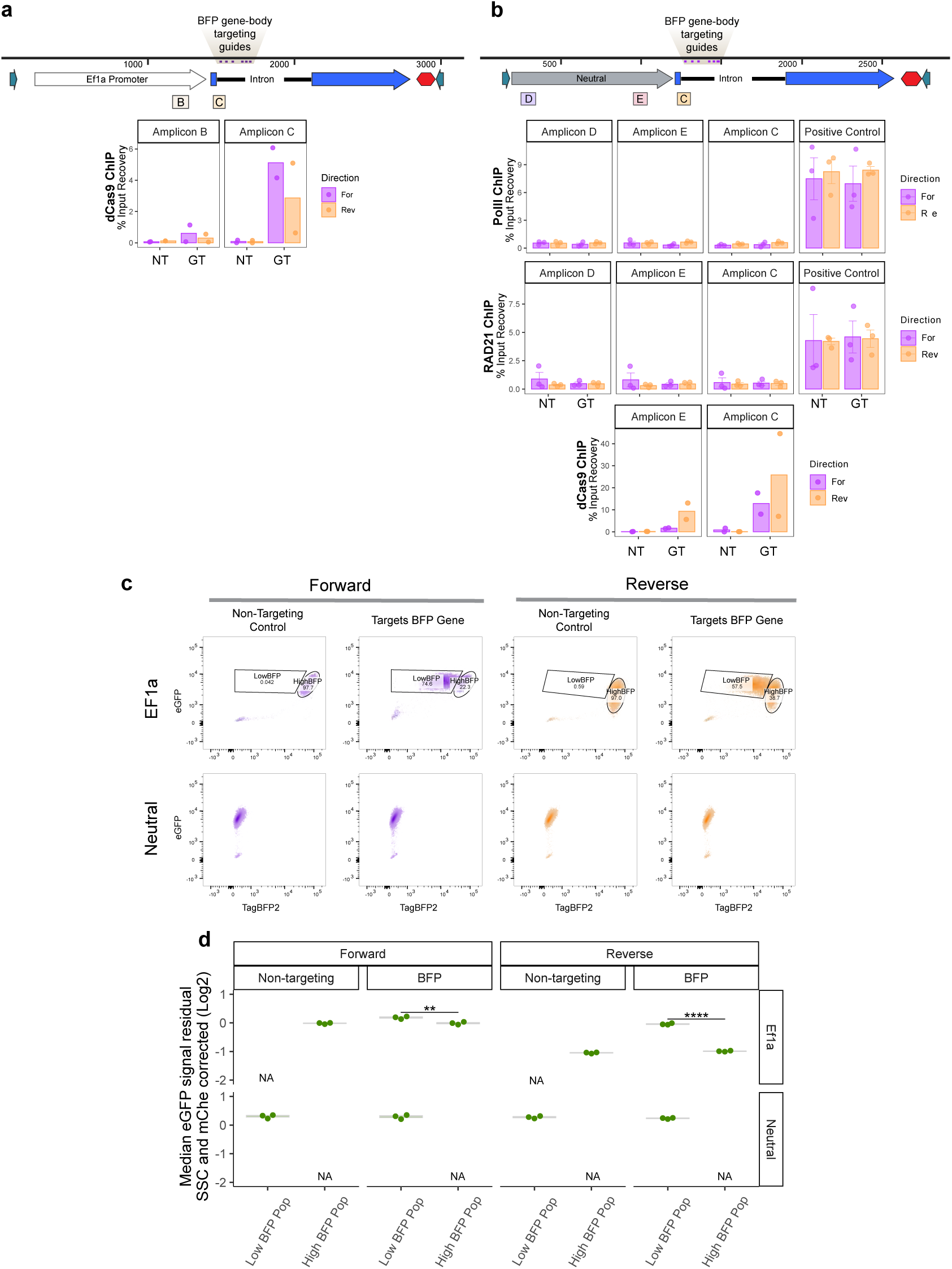
dCas9 CARGO controls in neutral insert line. **a)** dCas9 ChIP-qPCR percent input recovery at highlighted amplicon sites. n=2, bars represent means, NT=Non-targeting control, GT=Gene targeting CARGO. **b)** RNA Pol II, Rad21 and dCas9 ChIP-qPCR percent input recovery at highlighted amplicon sites in neutral integration lines. n=2 for dCas9, n=3 for Pol II and Rad21, mean ± s.e is plotted.**c)** Flow cytometry data showing “Low BFP” and “High BFP” gates in NT and GT populations for both Ef1a and neutral insert lines from representative replicate. All data from neutral integrations are considered “Low BFP”. **d)** Corrected median eGFP measurements from “Low BFP” and “High BFP” gates in non-targeting and gene-targeting populations, for both Ef1a and neutral insert lines. Median of measurements from 3 replicates plotted. Boxes represent the median, IQR, and whiskers (±1.5 IQR). Two-sided pairwise t-tests with Bonferroni correction, ****Padj ≤ 0.0001.

**Extended Data Fig. 7.**
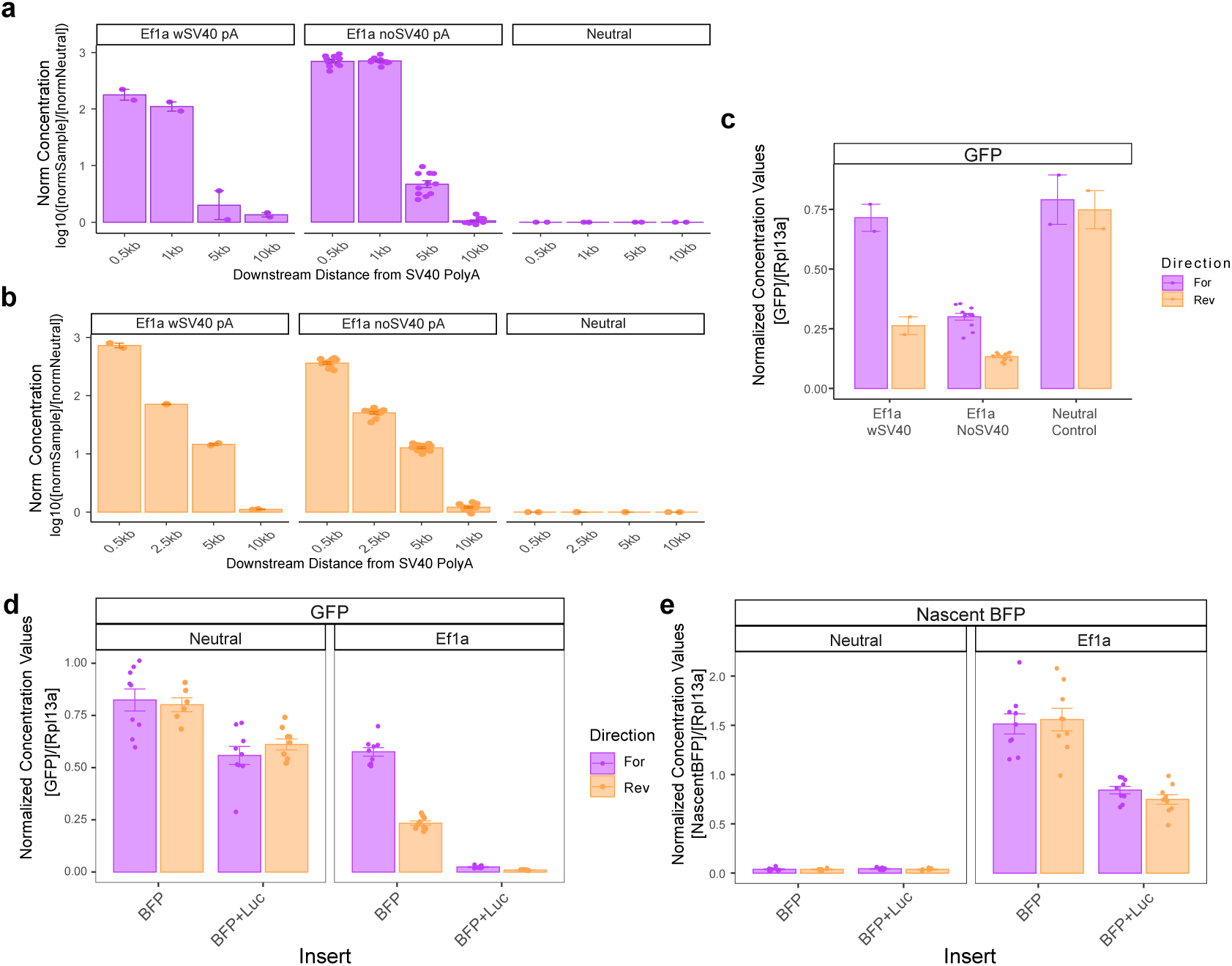
Confirmation of read-through from the integrated promoter and effects of lengthening the transcript on RNA levels. **a)** RT-PCR analysis of read-through transcript levels in forward Ef1a inserts. Absolute concentration of downstream transcripts was normalized to housekeeping Rpl13a, then normalized to concentration from neutral insert. 6 clones without SV40 polyA compared to 1 clone with Ef1a polyA and neutral polyA, in duplicate. mean ± s.e is plotted in log scale. **b)** RT-PCR analysis of read-through transcript levels in reverse Ef1a inserts. Absolute concentration of downstream transcripts was normalized to housekeeping Rpl13a, then normalized to concentration from neutral insert. 6 clones without SV40 polyA compared to 1 clone with Ef1a polyA and neutral polyA, in duplicate. mean ± s.e is plotted in log scale. **c)** RT-PCR analysis of eGFP transcript levels with or without SV40 polyA signal. Absolute concentration of eGFP transcripts normalized to housekeeping Rpl13a. 6 clones without SV40 polyA compared to 1 clone with Ef1a polyA and neutral polyA, in duplicate. Mean ± s.e is plotted. **d)** RT-PCR analysis of eGFP transcript levels in BFP and BFP+Luc insert lines. Absolute concentration of eGFP transcripts normalized to housekeeping Rpl13a. 3 biological replicates (independent lines) assayed three times. Mean ± s.e is plotted. **e)** RT-PCR analysis of nascent mTagBFP2 transcript levels in BFP and BFP+Luc insert lines. Absolute concentration of transcripts normalized to housekeeping Rpl13a. 3 biological replicates (independent lines) assayed three times. Mean ± s.e is plotted.

## Notes

https://www.ncbi.nlm.nih.gov/geo/query/acc.cgi?acc=GSE296229

https://www.ncbi.nlm.nih.gov/geo/query/acc.cgi?acc=GSE312207

### Competing Interest Statement

The authors have declared no competing interest.

### Summary of Updates

We added new experiments, updated figures, and text for this manuscript.

